## Supplementary Information for "Cell-free biosynthesis combined with deep learning accelerates de novo-development of antimicrobial peptides"

The supplementary information contains supplementary notes, tables, figures, and references:

Supplementary Notes 1-2 (p. 4-5)

**Supplementary Note 1:** Computational analysis of translation initiation rates.

**Supplementary Note 2:** Sequence similarity analyses using BLAST.

**Supplementary Note 3:** Molecular dynamics simulation of AMP-membrane interactions

Supplementary Tables 1-13 (p. 6-13)

**Supplementary Table 1:** Deep learning models training metrics.

**Supplementary Table 2:** Rounds of AMP generation, ranking, and testing with approaches and numbers.

**Supplementary Table 3:** Frequent 3- and 4-mers in non-AMPs, generated, prioritized, and tested and function AMPs.

**Supplementary Table 4:** Blast results of 30 functional AMPs against UniProt.

**Supplementary Table 5:** Blast results of 30 functional AMPs against the AMPs training dataset.

**Supplementary Table 6:** Blast results of 30 functional AMPs against 500 tested AMPs.

**Supplementary Table 7:** MIC against *E. coli* and *B.subtilis* and HC50 (hemolysis) and CC50 (cytotoxicity) value of AMPs on human red blood cells and HCT116 human colon cells, respectively.

**Supplementary Table 8:** MIC (minimum inhibitory concentration), HC50 (hemolysis), and CC50 (cytotoxicity) values of BP100 and Cecropin B measured in this study.

**Supplementary Table 9:** Membrane equilibration scheme.

**Supplementary Table 10:** Human plasma membrane composition.

**Supplementary Table 11:** *E. coli* inner membrane composition.

**Supplementary Table 12:** TFA-based cleavage cocktails.

**Supplementary Table 13:** Columns for analytical and (semi-)preparative HPLC-MS.

Supplementary Figures 1-39 (p. 14-32)

**Supplementary Fig. 1:** Growth curves in Fig. 2b with error bars.

**Supplementary Fig. 2:** SDS-PAGE gel images of AMPs produced using cell-free protein synthesis (CFPS).

**Supplementary Fig. 3:** RBS calculator results on the expressibility of AMPs.

**Supplementary Fig. 4:** Physicochemical properties and amino acid composition of AMPs.

**Supplementary Fig. 5:** Properties of two VAE models used in this study.

**Supplementary Fig. 6:** Complementary results on molecular dynamics simulations.

**Supplementary Fig. 7:** 21 days daily MIC measurements of the shortlisted AMPs for resistance study.

**Supplementary Figs. 8-34:** HPLC chromatogram of purified peptides.

Supplementary references (p. 33)

### Supplementary Notes:

**Supplementary Note 1:** Computational analysis of translation initiation rates.

Since de novo-designed AMPs have very diverse sequences, their translation can be greatly affected by mRNA folding[1] (**Supplementary Fig. 3a**). To examine the effect of mRNA sequences on translation, we used the RBS calculator[2] to predict the translation initiation rate (TIR) for each of the 500 tested AMPs. Despite AMPs being translated from the same RBS, the calculated TIR values are distributed in a wide range over four orders of magnitude (**Supplementary Fig. 3b**). Interestingly, the TIR values of the 30 functional AMPs are similarly distributed (**Supplementary Fig. 3b**) indicating that the translation initiation rate has not been a bottleneck in finding active AMPs. In addition to TIR which is calculated thermodynamically, the folding kinetic of mRNA also affects translation initiation such that the RBS calculator under-predicts slow folding mRNAs up to 10-fold[1]. Because mRNAs with low TIR fold slower (**Supplementary Fig. 3c**), their actual TIR value could be higher than the predicted one. This can narrow down the actual TIR range by an order of magnitude. Although we might have found more active AMPs if we had rationally designed RBS for all 500 tested AMPs, slow-translated functional peptides are more likely to have higher activity in killing bacteria in low amounts. These results show that cell-free production of AMPs enables the discovery of active AMPs despite their translation initiation rate, although the rational design of RBS for each AMP can lead to numerous functional AMPs.

**Supplementary Note 2:** Sequence similarity analyses using BLAST.
We sought to study the sequence similarities between our AMPs and both the training set and UniProt. For this purpose, we used the Basic Local Alignment Search Tool (BLAST) comparable to the previous works[3]. We assessed parameters including *E* (expected) value, percentage identity, query cover, and the raw alignment score. The *E* value is a parameter describing the number of hits that can be expected by chance in a dataset, also taking into account the length of a query. In BLAST searching for a query against the UniProt non-redundant database with ~240 million protein sequences, an *E* value ≤0.001 can refer to a significant match, hence homology, and the higher the *E* value, the higher the chance of coincidence. We performed BLAST for our 30 functional AMPs against the UniProt non-redundant database with an *E* value threshold of 10. For 25 AMPs we did not obtain any hit meaning that they had an *E* value >10. We saw a hit for only AMPs #6, #21, #23, #29, and #30 with *E* values of 7.1, 1.9, 9.7, 0.16, and 1.7, respectively (**Supplementary Table 4**). The BLAST hit with the lowest *E* value (0.16 for AMP #29 as the query) was a 511-amino acid bacterial outer membrane protein with an alignment score =40.5, percentage identity = 66.67%, and the query cover = 38%. Being a membrane protein and probably having a helical structure could be the reason for such a hit, nevertheless, the high *E* value and low percentage coverage in 38% of a peptide sequence do not indicate a homology. Additionally, none of the hits existed in the pretraining or training dataset. Next, we searched for sequence similarities between functional AMPs and the training dataset. Notably, inferring homology from the *E* value depends on the dataset size such that for the BLAST against our training dataset (~5000 sequences) the significance threshold would be around the *E* value of $10^{-7}$.[3] We obtained no significant hit (**Supplementary Table 5**), with the highest *E* value being 0.004 for a search between AMP #21 (GIGKFQKMRFIGAIRASKGVAKGLLRIAAIRTGRRALTT) and an AMP from the training dataset with the sequence of GILSTIKDFAIKAGKGAAKGLLEMASCKLSGQC. Moreover, BLAST searching of each functional AMP against all tested AMPs (including other functional ones) gave only one significant homology (**Supplementary Table 6**) meaning only one of the active AMPs

had a similar VAE-generated sequence (not a functional AMP). Altogether, these results demonstrate that the AMPs we found in this work are unique and diverse.

**Supplementary Note 3:** Molecular dynamics simulation of AMP-membrane interactions.

All selected 30 AMPs are characterized by a high proportion of basic, aromatic and hydrophobic residues. AlphaFold[4] predicts most of them as α-helical, either as one long helix or two shorter helices connected by a disordered turn (**Fig. 2b**). The only three exceptions are AMPs #10 (disordered), #8 and #14 (containing β-strands). Together, the structural prediction and their sequences suggest that most of the AMPs act as amphipathic peptides that preferably insert into the membrane interface region of negatively charged membranes, such as the inner membrane (IM) of bacteria. We performed molecular dynamics (MD) simulation of AMPs near models of the IM and the human plasma membrane (PM) **(Fig. 3a**). In our MD simulations all AMPs bound much stronger to the IM than to the PM. This is well reflected by the distributions of the distances of the centers of mass of the peptides to the membrane midplane (**Fig. 3b**). The distributions are much narrower and closer to the membrane core for the IM simulations than for the PM simulations.

In the MD simulations, the AMPs bound the IM within at most 200 ns after being released near the membrane (**Supplementary Fig. 6a**) and did not unbind again. On the PM on the other hand, AMP binding was only ever observed transiently. In no case did an AMP insert into the PM deeper than into the IM. This applies not only at the centers of mass of the complete peptides, but also at the level of any individual atom (**Supplementary Fig. 6b**). The preference for IM over PM binding is unlikely to be due to the tighter packing of the PM alone, as even high lateral membrane tension did not result in PM binding as close and as strong as IM binding without any imposed lateral tension. In general, however, the lateral tension allows tighter binding to either membrane compared with the simulations without lateral tension. Yet, even with high lateral tension, no AMPs spontaneously traversed either membrane in any of our simulations.

Furthermore, all AMPs have a higher number of interactions with the IM than with the PM (**Supplementary Fig. 6c**) and on the IM many of these contacts are electrostatic interactions between the basic peptides and the acidic phospholipid headgroups. These electrostatic interactions with the lipid headgroups on top of possible hydrophobic interactions with the lipid tails may explain why IM binding was irreversible on the 1 µs timescale of the MD simulations, whereas PM binding exhibited frequent un- and rebinding.

In our MD simulations, we observed that most AMPs did not fully retain their predicted mostly α-helical structure, but became more disordered as time progressed (**Supplementary Fig. 6d**). This partial unfolding was more pronounced for the AMPs in the PM systems, where the peptides spent more of the simulated time in the bulk solvent rather than at the membrane interface. This hints towards a general mechanism where the AMPs are unfolded in solution and adopt an ordered structure when bound to the membrane. The amphipathic character of the structured state is stabilized by membrane interactions. Altogether, these results imply that (i) these peptides are most likely to act on membranes and (ii) they prefer bacterial over human membranes.

#### Supplementary Tables:

**Supplementary Table 1:** models training metrics.

| **Regressor** | **Negative data*** | **Positive data*** | **Accuracy** | | |
|---|---|---|---|---|---|
| CNN_MIC_regressor_v0 | nonAMP[5] (5,582) | GRAMPA (5,102) | 0.786 | | |
| CNN_MIC_regressor_v1 | nonAMP[5] (5,582) | GRAMPA gram-specific 4,089 gram+, 4,619 gram- | 0.835 gram- 0.887 gram+ | | |
| CNN_MIC_regressor_v2 | UniProtKB nonAMP 10,612 | GRAMPA gram-specific 4,089 gram+, 4,619 gram- | 0.932 gram- 0.947 gram+ | | |
| RNN_MIC_regressor_v0 | UniProtKB nonAMP 10,612 | GRAMPA gram-specific 4,089 gram+, 4,619 gram- | 0.947 gram- 0.942 gram+ | | |
| CNN_tox_classifier | 17,434 | 8,992 | 0.942 | | |
| **Generator** | **Pretraining data** | **Training data** | **KL** term** | **KL loss** | **Recon. loss** |
| VAE_v0 | - | GRAMPA (5,319) | - | 1.179 | 3.331 |
| VAE_v1 | UniProtKB (~1.5 M) | GRAMPA (5,319) | - | 1.265 | 3.027 |
| VAE_v2 | UniProtKB (~1.5 M) | GRAMPA (5,319) | + | 2.494 | 4.925 |

*See Methods. ** Kullback-Leibner.

**Supplementary Table 2:** Rounds of AMP generation, filtering, and prioritization with different models/approaches and numbers.

| | **round 0** | **round 1** | **round 2** | **round 3** | **round 4** |
|---|---|---|---|---|---|
| **Generator** | VAE_v0 | VAE_v0 | VAE_v1 | VAE_v1 | VAE_v2 |
| **Sampling** | Optimized Cecropin B* | random | random | random | random |
| **Regressor** | CNN reg_v0 | CNN reg._v0 | CNN reg._v1 | CNN reg._v2 + Toxicity classifier | CNN reg._v2 + RNN reg._v2 |
| **All generated peptides** | 100 | 100,000 | 100,000 | 200,000 | 150,000 |
| **Viable peptides** | 100 | 9,220 | 9,117 | 18,218 | 29,457 |
| **MIC-predicted and experimentally tested** | 50 | 50 | 150 | 100 | 150 |
| **Functional peptides discovered** | 0 | 2 | 9 | 0 | 19 |
| **Efficiency (%)** | **0** | **4** | **6** | **0** | **12.6** |

For simplicity, in Fig. 2c we included the two functional AMPs from round 1 into the latent space of VAE_v1 together with functional AMPs of round 2. VAE_v0 had the same architecture and loss function as VAE_v1 however it was not pretrained. *Gradient descent optimization at the neighborhood of Cecropin B in the latent space.

**Supplementary Table 3:** Frequent 3- and 4-mers in non-AMPs, generated, prioritized and tested and function AMPs.

| k-mers | UniProt nonAMP | Frequency | VAE training AMPs | Frequency | 500 test AMPs | Frequency | 30 functional AMPs | Frequency |
|---|---|---|---|---|---|---|---|---|
| 3-mers | RRR | 0.0046 | LKK | 0.0061 | RRR | 0.0125 | KKK | 0.0172 |
| | KRT | 0.0026 | KKL | 0.0053 | KKK | 0.0118 | FKK | 0.0097 |
| | KGR | 0.0025 | LLK | 0.0051 | FKK | 0.0056 | KKF | 0.0075 |
| | GRK | 0.0024 | KLL | 0.0045 | FFF | 0.0056 | FFK | 0.0067 |
| | RKR | 0.0024 | KKI | 0.0044 | KKG | 0.0042 | KKG | 0.006 |
| | MKV | 0.0022 | AKK | 0.0034 | LLL | 0.0032 | LFL | 0.0052 |
| | RQG | 0.0022 | PRP | 0.0031 | RKK | 0.003 | FKA | 0.0045 |
| | VLA | 0.0022 | AGK | 0.0031 | GFK | 0.0028 | FFF | 0.0045 |
| | LAV | 0.0022 | LAK | 0.003 | FLF | 0.0028 | RRR | 0.0045 |
| | GFR | 0.0022 | AAK | 0.003 | KKF | 0.0026 | KKY | 0.0037 |
| | VIC | 0.0022 | IKK | 0.0029 | FFK | 0.0026 | GRR | 0.0037 |
| | GLL | 0.0021 | GLL | 0.0028 | RRL | 0.0026 | RRF | 0.0037 |
| 4-mers | KQRQ | 0.0019 | KLLK | 0.0022 | RRRR | 0.0055 | KKKK | 0.0061 |
| | HGFR | 0.0018 | LKKL | 0.002 | KKKK | 0.0041 | FKKK | 0.0038 |
| | QRQG | 0.0018 | LLKK | 0.0018 | FFFF | 0.0017 | FKAR | 0.0023 |
| | HKQR | 0.0018 | KKLL | 0.0018 | FKKK | 0.0015 | KKFV | 0.0023 |
| | MKRT | 0.0016 | ASKV | 0.0013 | KKGF | 0.0012 | FFFF | 0.0023 |
| | MKVR | 0.0015 | RPRP | 0.0012 | KKKG | 0.001 | FFKK | 0.0023 |
| | RRRR | 0.0014 | KKKK | 0.0012 | FFKK | 0.001 | KKKY | 0.0023 |
| | KHKQ | 0.0013 | LKKI | 0.0012 | KQKK | 0.001 | FCFK | 0.0023 |

**Supplementary Table 4:** NCBI BLASTP 2.13.0+ results of 30 functional AMPs against nonredundant UniProt containing 498M (498,091,743) sequences on August 5, 2022. The word size of 6, expect threshold of 10, PAM30 matrix, gap initiation penalty of 9 and gap extension penalty of 1, conditional compositional score matrix adjustment, and low complexity regions filter were used. The AMPs not shown here did not return any hit with an *E* value threshold of 10.

| AMP | Hit reference ID | *E* value | Percent Identity | Query Cover | Max Score | Bit Score |
|---|---|---|---|---|---|---|
| AMP #6 | MBW0556277.1 | 7.1 | 85.71% | 28% | 38.4 | 38.4 |
| AMP #21 | MCB5272487.1 | 1.9 | 61.54% | 65% | 39.2 | 39.2 |
| AMP #23 | REE03804.1 | 9.7 | 60.00% | 55% | 37.5 | 37.5 |
| AMP #29 | MBR6433148.1 | 0.16 | 59.26% | 45% | 43.5 | 43.5 |
| AMP #30 | ORE05204.1 | 1.7 | 66.67% | 38% | 40.5 | 40.5 |

**Supplementary Table 5:** NCBI BLASTP 2.13.0+ results of 30 functional AMPs against the training dataset containing ~5000 AMPs. The word size of 2, expect threshold of 10, BLOSUM62 matrix, gap initiation penalty of 11 and gap extension penalty of 1, and conditional compositional score matrix adjustment were used. The AMPs not shown here did not return any hit with an *E* value threshold of 10. Note that inferring homology from the *E* value depends on the dataset size such that for the BLAST against the training dataset the significance threshold would be around the *E* value of $10^{-7}$.[3]

| AMP | Hit | Bit score | *E* value | AMP | Hit | Bit score | *E*-value |
|---|---|---|---|---|---|---|---|
| AMP #1 | train1855<br>train3986<br>train5028 | 16.9<br>16.9<br>16.9 | 7.7<br>8.4<br>8.8 | AMP #16 | train1491<br>train3557<br>train75<br>train2471<br>train5194<br>train3400<br>train1334 | 20.0<br>17.7<br>17.7<br>17.3<br>16.5<br>16.5<br>16.5 | 0.41<br>2.8<br>3.4<br>4.3<br>8.5<br>8.7<br>9.8 |
| AMP #2 | train68<br>train941<br>train3543<br>train939<br>train952<br>train579<br>train248 | 18.9<br>18.1<br>18.1<br>17.7<br>17.3<br>17.3<br>17.3 | 1.8<br>2.9<br>2.9<br>3.0<br>5.0<br>5.3<br>6.9 | AMP #17 | train471<br>train3918<br>train2948 | 18.5<br>18.1<br>16.9 | 1.6<br>2.7<br>7.5 |
| AMP #3 | train5167<br>train1437<br>train1426<br>train1425 | 17.3<br>16.9<br>16.9<br>16.9 | 4.4<br>8.2<br>8.6<br>9.1 | AMP #18 | train3793<br>train2453 | 17.3<br>16.9 | 6.2<br>7.8 |

| | | | | | | | |
|---|---|---|---|---|---|---|---|
| AMP #5 | train3915<br>train3957<br>train3528<br>train2105 | 18.1<br>16.9<br>16.5<br>16.9 | 2.4<br>5.7<br>8.2<br>8.5 | AMP #19 | train4740<br>train2912<br>train2509<br>train1446<br>train1447<br>train1315 | 19.6<br>19.6<br>18.1<br>17.7<br>17.7<br>16.9 | 0.70<br>0.74<br>2.6<br>4.9<br>5.0<br>9.0 |
| AMP #6 | train1503<br>train4288<br>train1185<br>train707 | 19.6<br>16.9<br>16.5<br>16.9 | 0.67<br>7.7<br>8.5<br>9.0 | AMP #21 | train1667<br>train1612<br>train1611<br>train2133<br>train2035<br>train2135<br>train2036 | 25.0<br>17.7<br>17.7<br>17.7<br>17.7<br>16.5<br>16.5 | **0.004***<br>2.8<br>3.0<br>3.0<br>3.1<br>7.7<br>9.4 |
| AMP #7 | train4365 | 16.9 | 9.2 | AMP #22 | train529<br>train832<br>train1181<br>train833<br>train2604 | 18.1<br>17.3<br>16.5<br>16.5<br>16.2 | 2.5<br>4.0<br>7.3<br>7.7<br>9.8 |
| AMP #8 | train4630<br>train383<br>train2130 | 17.7<br>17.3<br>16.9 | 3.8<br>7.2<br>9.4 | AMP #23 | train1564<br>train1563 | 18.5<br>18.1 | 1.4<br>2.2 |
| AMP #10 | train3299<br>train746<br>train3300<br>train744<br>train4741<br>train543 | 22.3<br>19.6<br>18.5<br>17.7<br>17.3<br>16.5 | 0.06<br>0.49<br>2.4<br>3.2<br>7.9<br>8.3 | AMP #24 | train1932<br>train1931<br>train2063<br>train87<br>train4217 | 22.3<br>20.0<br>18.5<br>16.9<br>16.9 | 0.095<br>0.45<br>1.7<br>7.0<br>8.1 |
| AMP #11 | train3881 | 21.6 | 0.18 | AMP #25 | train4867<br>train2475<br>train2474<br>train469 | 18.5<br>17.7<br>17.7<br>17.3, | 1.7<br>2.9<br>2.9<br>5.9 |
| AMP #12 | train2472 | 17.3 | 9.7 | AMP #27 | train1705<br>train4437 | 17.7<br>16.5 | 4.0<br>7.7 |
| AMP #13 | train1301<br>train4915<br>train1881 | 17.7<br>16.9<br>16.5 | 4.0<br>6.5<br>10.0 | AMP #28 | train1263 | 17.7 | 2.9 |
| AMP #14 | train9 | 18.1 | 2.9 | AMP #29 | train1368 | 16.9 | 7.6 |
| AMP #15 | train3476<br>train63<br>train1746<br>train3475 | 18.5<br>18.1<br>16.9<br>16.5 | 2.0<br>2.8<br>8.2<br>8.2 | AMP #30 | train1298<br>train831<br>train4893<br>train1297 | 19.2<br>18.1<br>17.7<br>16.9 | 1.1<br>2.6<br>4.9<br>8.0 |

* Lowest E-value (not sinificant): AMP #21 (GIGKFQKMRFIGAIRASKGVAKGLLRIAAIRTGRRALTT) vs train1667 (GILSTIKDFAIKAGKGAAKGLLEMASCKLSGQC)

**Supplementary Table 6:** NCBI BLASTP 2.13.0+ results of 30 functional AMPs against 500 tested AMPs. The word size of 2, expect threshold of 10, BLOSUM62 matrix, gap initiation penalty of 11 and gap extension penalty of 1, and conditional compositional score matrix adjustment were used. Note that inferring homology from the *E* value depends on the dataset size such that for the BLAST against the training dataset the significance threshold would be around the *E* value of $10^{-8}$.[3] All hits with an *E* value <10 are provided as a **Source Data** file.

| ***E* value** | **>10** | **1-10** | **$10^{-3}$-1** | **$10^{-8}$-0.$10^{-3}$** | **<0.$10^{-8}$** |
|---|---|---|---|---|---|
| Count | 14782 | 175 | 41 | 1 | 1* |

*The only significant *E*-value (1.00E-16): AMP #30 (GFGLWGLFHFKMNVPNLFKNGFIFLIIMIFTVWGLFFGKKKAYIEKFL) vs gen66 (GFGLWLLFQFKIRPPRLFKNGFLFLILMIFTTWILFFVKQKLFGMPFL)

**Supplementary Table 7:** MIC, HC50 and CC50 values (µM) of the plot in **Fig. 4a**. The values are the average of n=3 and n = 2 independent experiments for MIC and HC50/CC50 respectively.

| | ***E. coli* MIC** | ***B. subtilis* MIC** | **CC50** | **HC50** |
|---|---|---|---|---|
| AMP #1 | >100 | 0.8 | 113.0 | >250 |
| AMP #3 | 12.5 | 0.8 | 68.0 | >250 |
| AMP #5 | 2.1 | 0.5 | 68.4 | 82.9 |
| AMP #6 | 25.0 | 2.1 | 67.5 | 73.8 |
| AMP #7 | >100 | 1.6 | 132.9 | >250 |
| AMP #9 | 25.0 | 3.1 | 146.0 | >250 |
| AMP #10 | 50.0 | 0.6 | 105.0 | >250 |
| AMP #12 | 37.5 | 6.3 | >250 | 17.1 |
| AMP #13 | 25.0 | 1.6 | 25.8 | >250 |
| AMP #14 | 37.5 | 6.3 | >250 | >250 |
| AMP #15 | 25.0 | 0.8 | 30.7 | 24.8 |
| AMP #16 | 6.3 | 0.8 | 39.3 | 180.2 |
| AMP #17 | 12.5 | 6.3 | 133.7 | >250 |
| AMP #18 | 50.0 | 3.1 | 153.0 | 4.7 |
| AMP #19 | 25.0 | 8.4 | >250 | >250 |
| AMP #21 | 10.4 | 0.5 | >250 | >250 |
| AMP #23 | 20.8 | 1.6 | 79.1 | 11.5 |
| AMP #24 | 25.0 | 1.6 | 91.7 | >250 |
| AMP #26 | 100.0 | 37.5 | >250 | >250 |
| AMP #27 | 12.5 | 0.4 | 75.7 | 105.1 |
| AMP #28 | 25.0 | 0.4 | 145.0 | >250 |
| AMP #29 | 12.5 | 6.3 | >250 | 50.6 |

**Supplementary Table 8:** MIC (minimum inhibitory concentration), HC50 (hemolysis), and CC50 (cytotoxicity) values of BP100 and Cecropin B measured in this study. The values are the average of n=3 independent experiments for *E. coli* and *B. subtilis* MIC and n = 2 independent experiments for others (n.d., not detected).

| MIC/HC50/CC50 (µM) | PB100 | Cecropin B |
|---|---|---|
| *Escherichia coli* MIC | 3.1 | 0.4 |
| *Bacillus subtilis* MIC | 0.5 | 2.6 |
| *Acinetobacter baumannii* MIC | 0.6 | 0.2 |
| *Enterobacter cloacae* MIC | 12.5 | 0.4 |
| *Klebsiella pneumoniae* MIC | 2.4 | 0.2 |
| *Pseudomonas aeruginosa* MIC | 6.3 | 3.1 |
| *Staphylococcus aureus (MRSA)* MIC | 6.3 | >25 |
| *Enterococcus faecium* MIC | 4.7 | >25 |
| Hemolysis (HC50) | 96.0 | nd |
| Cytotoxicity (CC50) | 56.8 | 154.2 |

**Supplementary Table 9:** Membrane equilibration scheme.

| Step | Time[ns] | Timestep [fs] | Ensemble | Headgroup position restraints [kJ mol$^{-1}$] | Tail dihedral angle restraints [kJ mol$^{-1}$] |
|---|---|---|---|---|---|
| **EM** | | | | 1000 | 1000 |
| **1** | 1.25 | 1 | NVT | 1000 | 1000 |
| **2** | 1.25 | 1 | NVT | 400 | 400 |
| **3** | 1.25 | 1 | NPT | 400 | 200 |
| **4** | 0.5 | 2 | NPT | 200 | 200 |
| **5** | 0.5 | 2 | NPT | 40 | 100 |
| **6** | 0.5 | 2 | NPT | 0 | 0 |

**Supplementary Table 10:** Human plasma membrane composition.

| Lipid | Full name | Abundance [%] |
|---|---|---|
| CHOL | Cholesterol | 35.8 |
| PSM | N-palmitoyl-D-erythro-sphingosylphosphorylcholine | 13.1 |
| NSM | N-nervonoyl-D-oleoyl-sphingosylphosphorylcholine | 10.0 |
| LSM | N-lignoceroyl-D-oleoyl-sphingosylphosphorylcholine | 8.4 |
| PLPC | 1-palmitoyl-2-linoleoyl-sn-glycero-3-phosphatidylcholine | 16.2 |
| SOPC | 1-stearoyl-2-oleoylphosphatidylcholine | 7.5 |
| PAPC | 1-palmitoyl-2-arachidonoyl-glycero-3-phosphatidylcholine | 5.6 |
| PLA20(PE) | 1-O-stearoyl-2-O-arachidonoyl-glycero-3-phosphatidylethanolamine | 2.2 |
| SAPS | 1-stearoyl-2-arachidonoyl-glycero-3-phosphatidylserine | 1.2 |

**Supplementary Table 11:** *E. coli* inner membrane composition.

| Lipid | Full name | Abundance [%] |
|---|---|---|
| PVPE | 1-palmitoyl-2-vacenoyl-sn-glycero-3-phosphatidylethanolamine | 75 |
| PVPG | 1-palmitoyl-2-vacenoyl-sn-glycero-3-phosphatidylglycerol | 20 |
| PVCL2 | 1-palmitoyl-2-vacenoyl-cardiolipin | 5 |

**Supplementary Table 12:** TFA-based cleavage cocktails. Depending on the content of the oxidation prone amino acids Cys, Met and Trp one of the following cleavage cocktails has been used.

| Cocktail | Cocktail Composition (v/v) |
| --- | --- |
| Cleavage Cocktail A | 82.5% TFA, 5.0% $H_2O$, 5.0% phenol, 5.0% thioanisole, 2.5 % EDT |
| Cleavage Cocktail B | 90 % TFA, 4.0% TMSBr, 4.0% thioanisole, 2.0% EDT |

**Supplementary Table 13:** Columns for analytical and (semi-)preparative HPLC-MS. Column 1 was used for the preparative purification of peptide crude while column 2 was for the characterization of the final purified peptides.

| Column | Type | Dimensions | Flow | Purpose |
| --- | --- | --- | --- | --- |
| Column 1 | XBridge Prep C18 OBD (Waters) | 250 x 19 mm, 5 µm | 16 mL/min | purification |
| Column 2 | Eclipse XBD-C18 (Agilent Technologies) | 150 x 4.6 mm, 5 µm | 1 mL/min | analysis |

Supplementary Figures:

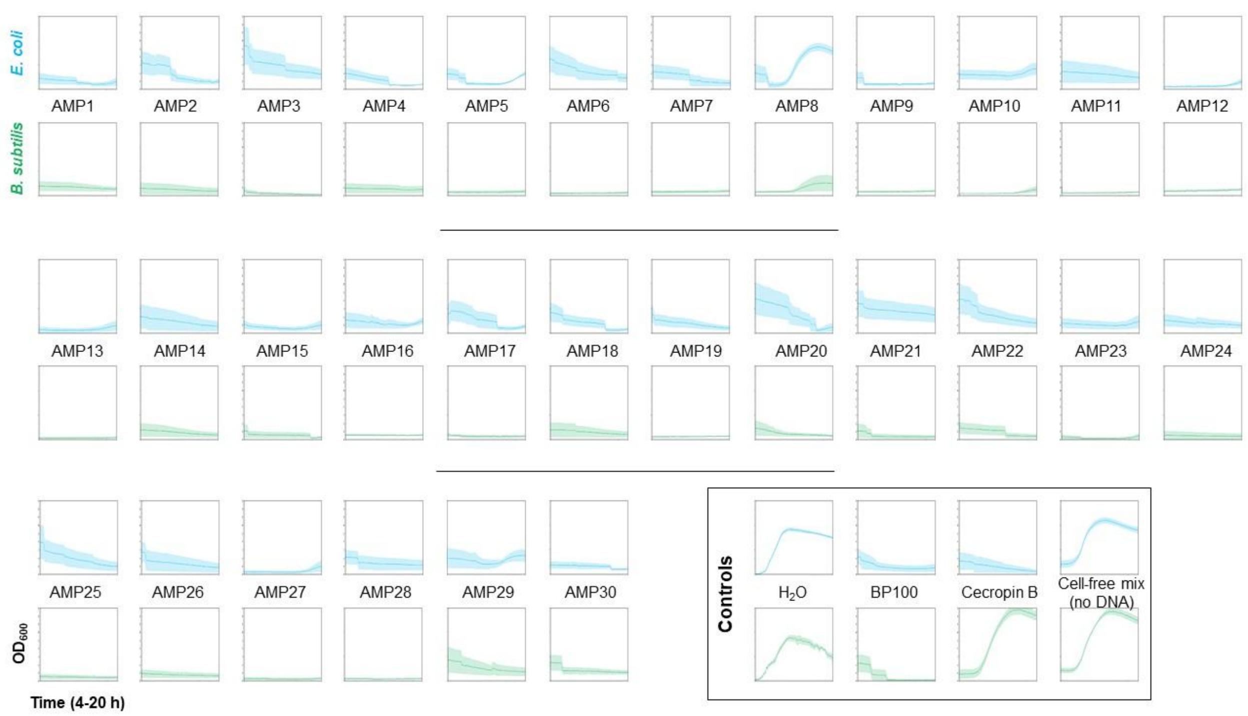

**Supplementary Fig. 1:** $OD_{600}$ over time 4-20 h growth curves (n=3 independent experiments) in **Fig. 2b** with error bars as standard deviation. Raw data is provided as a **Source Data** file.

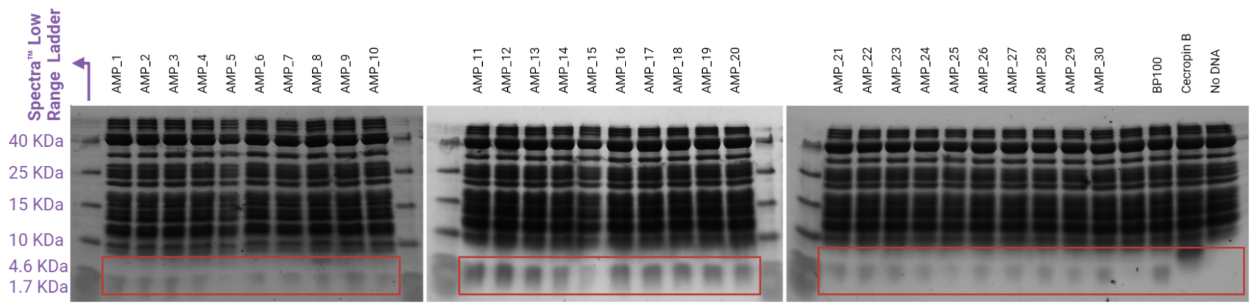

**Supplementary Fig. 2:** SDS-PAGE of AMPs produced using cell-free protein synthesis. This experiment was done with no replicates and the raw images are provided as a **Source Data** file.

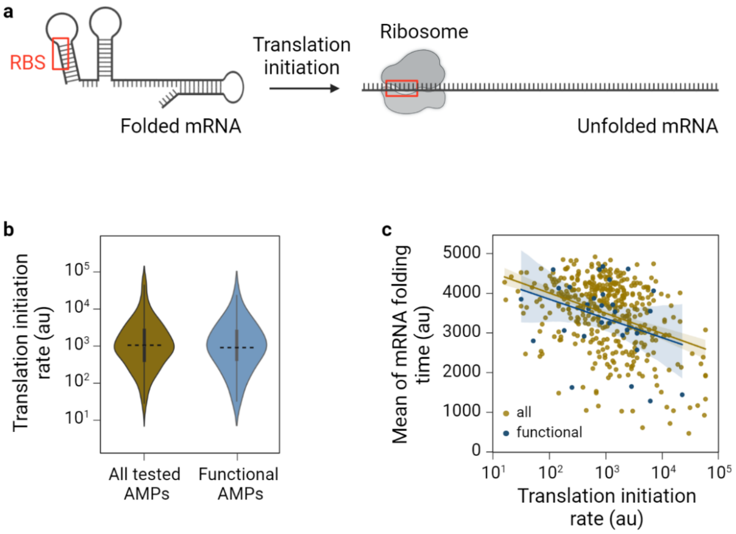

**Supplementary Fig. 3:** RBS calculator simulations on the translation of AMPs. **a,** Schematic of the translation initiation by mRNA unfolding and binding the ribosome to the RBS (ribosome binding site). **b,** The translation initiation rate (TIR) calculated using the RBS calculator for all 500 tested AMPs (yellow) and 30 functional AMPs (blue). **c,** Mean of mRNA folding time versus translation initiation rate for all 500 tested AMPs (yellow) and 30 functional AMPs (blue). RBS Calculator v1.0[2] was used. For folding times, we used Kinfold[6] run 1000 times for each sequence, with a folding time cut-off of 5000 and the rest of the parameters as default, EnergyModel: dangle=2 Temp=37.0 logML=logarithmic Par=VRNA-1.4, MoveSet: noShift=off noLP=off, Simulation: num=1000 time=5000.00 seed=clock fpt=on mc=Kawasaki, Simulation: phi=1 pbounds=0.1 0.1 2.

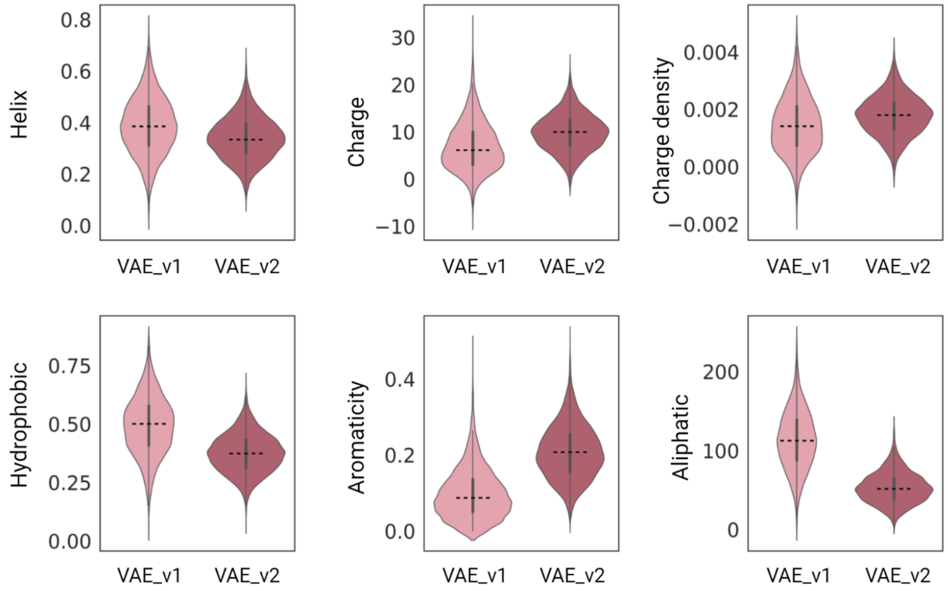

**Supplementary Fig. 4:** Properties of two VAE models used in this study. These features were calculated on viable peptides with 36-48 amino acids generated using each VAE. These features were computed using Biopython 1.79[7] and the modlAMP 4.3.0[8] packages.

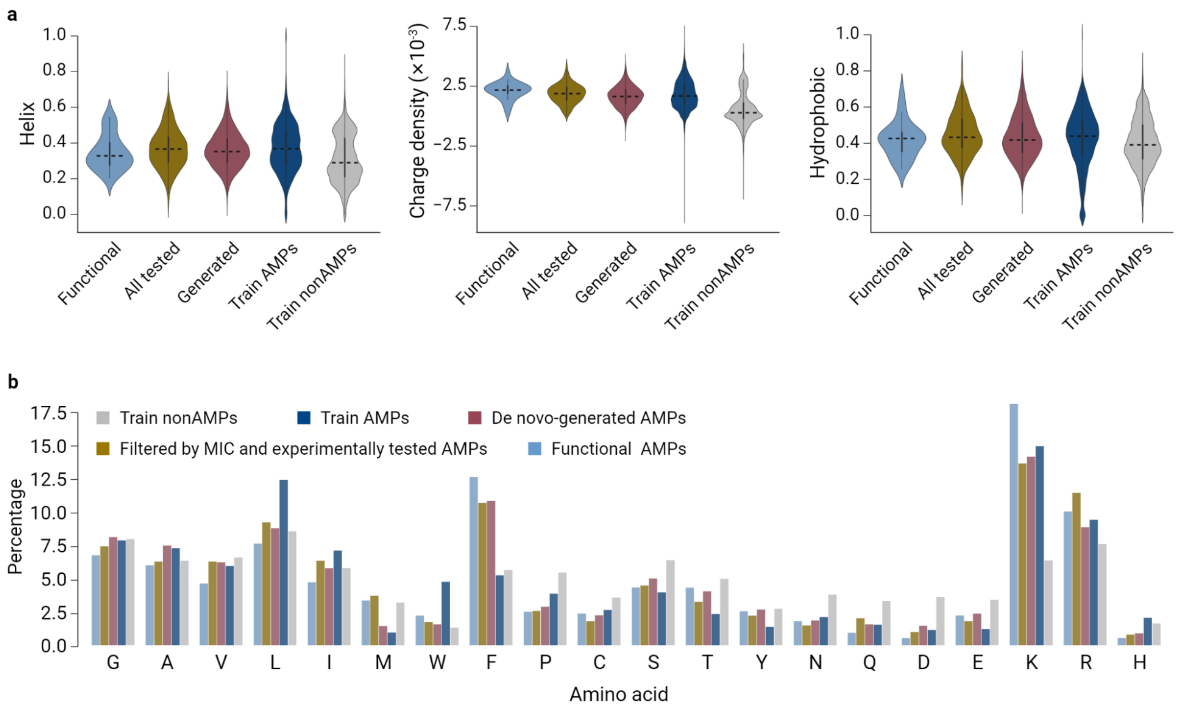

**Supplementary Fig. 5:** Physicochemical properties (**a**) and amino acid composition of AMPs (**b**) computed using Biopython 1.79[7] and the modlAMP 4.3.0[8] packages. Source data for (**b**) are provided as a **Source Data** file.

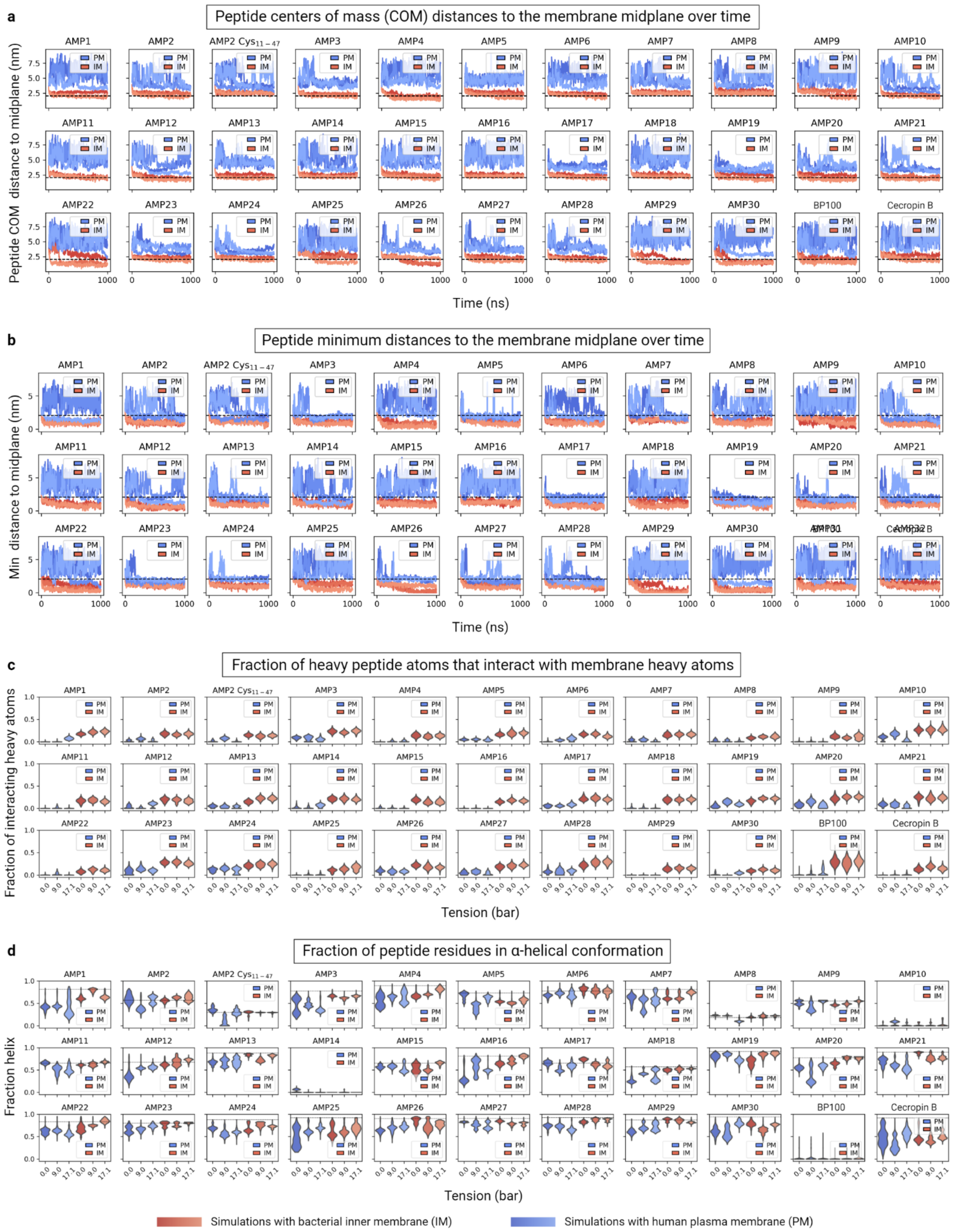

**Supplementary Fig. 6:** Complementary results on molecular dynamics simulations. **a,** Distances along the direction of the membrane normal between the AMP centers of mass and the membrane midplane plotted vs. time for each replicate, run with different lateral tensions (darkest: 0 bar; lighter: 9 bar; lightest: 17.1 bar) and with different membranes (blue: PM; red: IM). The dotted line indicates the headgroup phosphate positions. **b,** Minimum distances along the direction of the membrane normal between any heavy AMP atom and the membrane midplane plotted vs. time for each replicate run with different lateral tensions (darkest: 0 bar; lighter: 9 bar; lightest: 17.1 bar) and with different membranes (blue: PM; red: IM). The dotted line indicates the headgroup phosphate positions. **c,** Distributions of the fractions of heavy peptide atoms that interact with membrane heavy atoms (cutoff 3.5 Å). Distributions are calculated from the last 950 ns of 1 µs long replicates, run with different lateral tensions and with different membranes (blue: PM; red: IM). **d,** Distributions of the fractions of AMP residues that are in α-helical conformation. Distributions are calculated from the last 950 ns of 1 µs long replicates, run with different lateral tensions and with different membranes (blue: PM; red: IM). The dotted line indicates the helicity in the initial model prediction by AlphaFold[4].

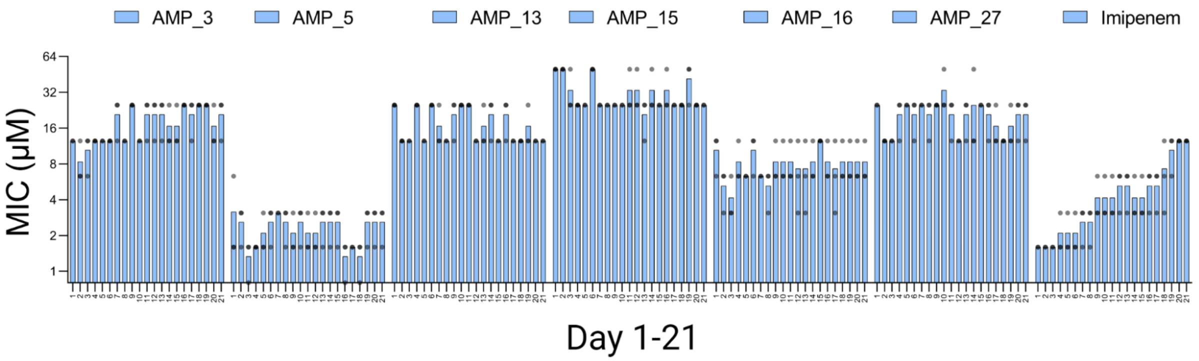

**Supplementary Fig. 7.** 21 days daily MIC measurements of the six broad-band AMPs for resistance study. The y axis is in log2 scale. See also **Fig 6a**. Bars are the average of n = 3 independent experiments. Source data for this figure are provided as a **Source Data** file.

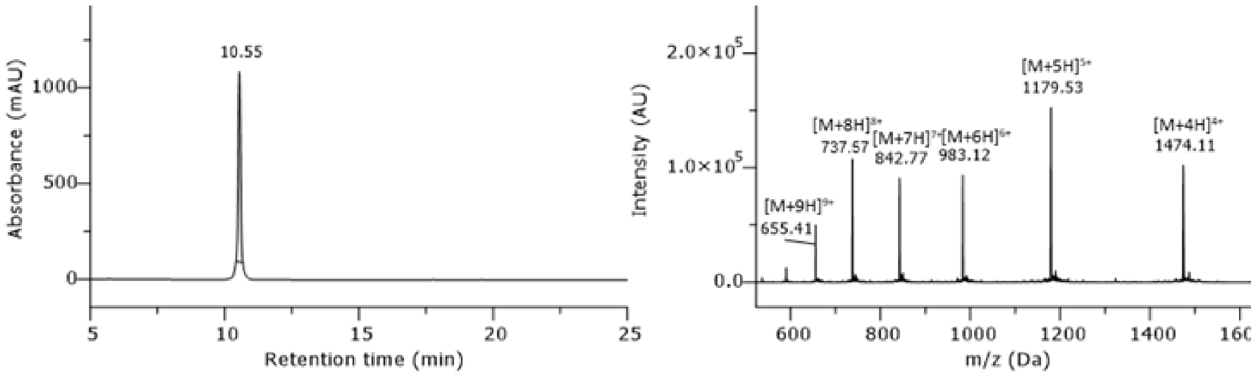

**Supplementary Fig. 8: HPLC chromatogram of purified peptide AMP #1.** Gradient 5-95% B in column 2, monitored at 220 nm.

$H_2N$-MLFGSRAKKYGKEAKQEKSFQYPKSSFVACKKKWKRSKHFFKTFKKKVS-$CONH_2$; peptide has been synthesized in 5 µmol scale. After purification (column 1, 05-50% MeCN), the 19 x TFA salt product (9.40 mg, 1.17 µmol, 23%) was obtained as a white solid. $t_R$ = 10.55 min. Purity ≥ 99%. Formula: $C_{275}H_{434}N_{76}O_{64}S_2$. Molecular weight: 5892.98 g/mol. HRMS-ESI+ (m/z): $[M+10H]^{10+}$ calcd.: 590.2329; found: 590.2328.

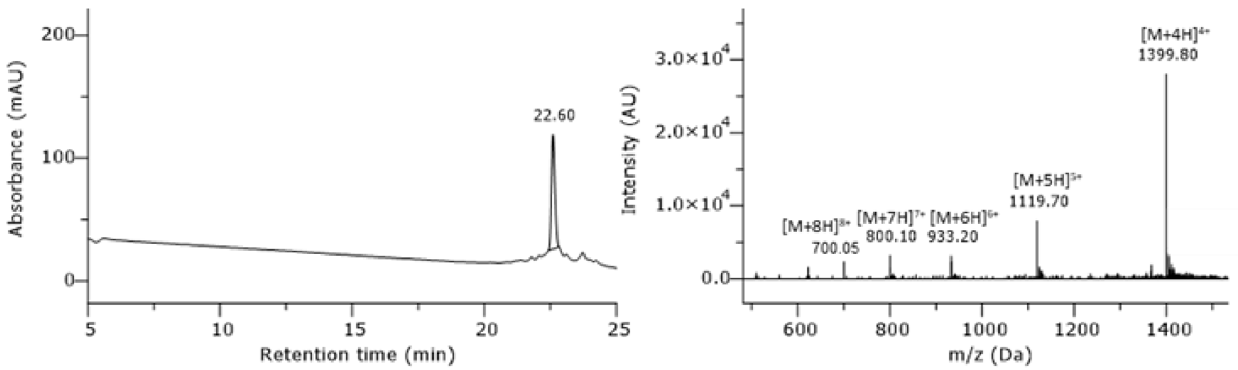

**Supplementary Fig. 9: HPLC chromatogram of purified peptide AMP #2**. Gradient 15-35% B, with addition of 1 mM TCEP, in column 2, monitored at 220 nm.

$H_2N$-MWLKMRKCCGCGFKYCLKCVQKKGRIFKTLGKAKMWPKWFFKIGGKC-$CONH_2$; peptide has been synthesized in 5 µmol scale. After purification (column 1, 05-50% MeCN), the 15 x TFA salt product (3.49 mg, 0.47 µmol, 9%) was obtained as a white solid. $t_R$ = 22.60 min. Purity > 83%. Formula: $C_{259}H_{412}N_{70}O_{50}S_9$. Molecular weight: 5595.06 g/mol. HRMS-ESI+ (m/z): $[M+9H]^{9+}$ calcd.: 622.5562; found: 622.5576.

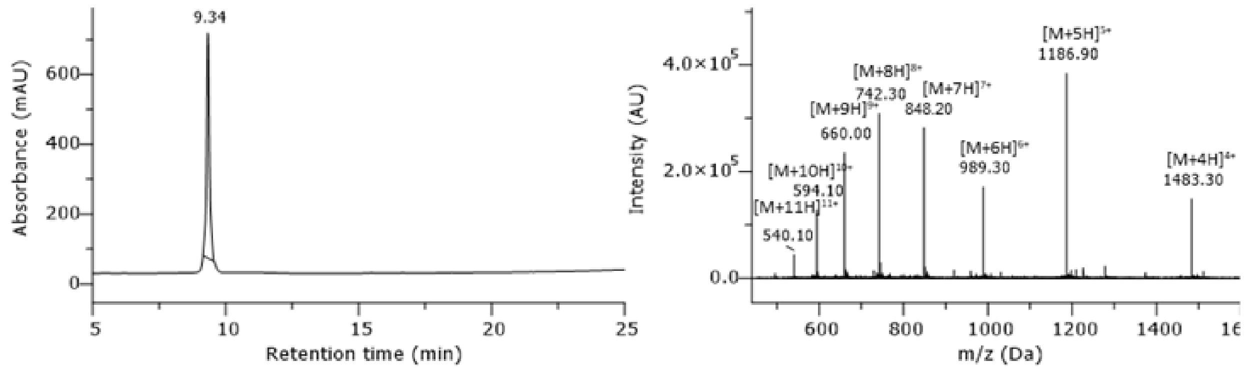

**Supplementary Fig. 10: HPLC chromatogram of purified peptide AMP #3.** Gradient 5-95% B in column 2, monitored at 220 nm.

$H_2N$-MNNRGPLGRRFKARRKWKKFVAGKMKKKRKRFKGFKKKGGFTPFVKKFV-$CONH_2$; peptide has been synthesized in 5 μmol scale. After purification (column 1, 05-35% MeCN), the 23 x TFA salt product (9.60 mg, 1.12 μmol, 22%) was obtained as a white solid. $t_R$ = 9.34 min. Purity ≥ 99%. Formula: $C_{277}H_{457}N_{89}O_{52}S_2$. Molecular weight: 5930.28 g/mol. HRMS-ESI+ (m/z): $[M+7H]^{7+}$ calcd.: 848.0840; found: 848.0827.

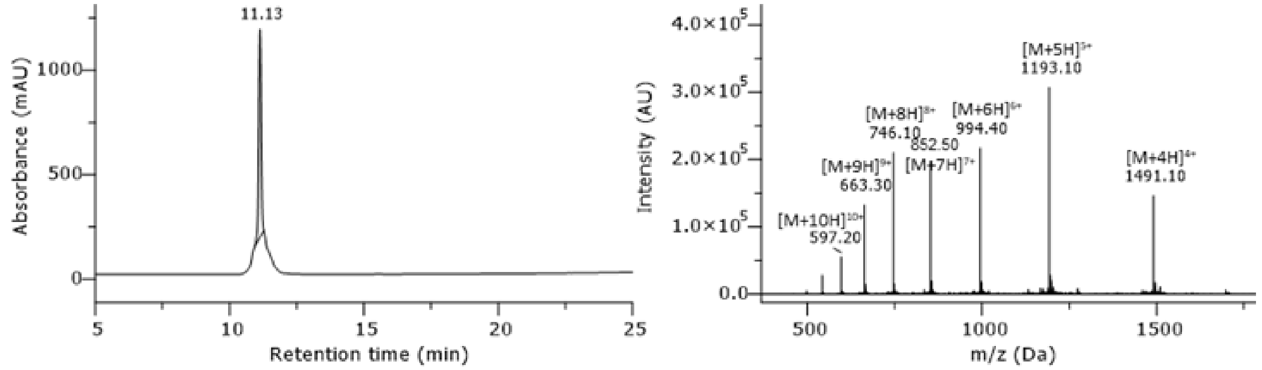

**Supplementary Fig. 11: HPLC chromatogram of purified peptide AMP #5.** Gradient 5-95% B in column 2, monitored at 220 nm.

$H_2N$-MRRKTPVKWKTFFKALKHKKKIFKKTFEKFKFLAKGPAFLKGFDQKLKS-$CONH_2$; peptide has been synthesized in 5 μmol scale. After purification (column 1, 05-50% MeCN), the 20 x TFA salt product (9.27 mg, 1.12 μmol, 22%) was obtained as a white solid. $t_R$ = 11.13 min. Approximate purity > 90%. Formula: $C_{289}H_{462}N_{76}O_{58}S$. Molecular weight: 5961.29 g/mol. HRMS-ESI+ (m/z): $[M+7H]^{7+}$ calcd.: 852.5122; found: 852.5127.

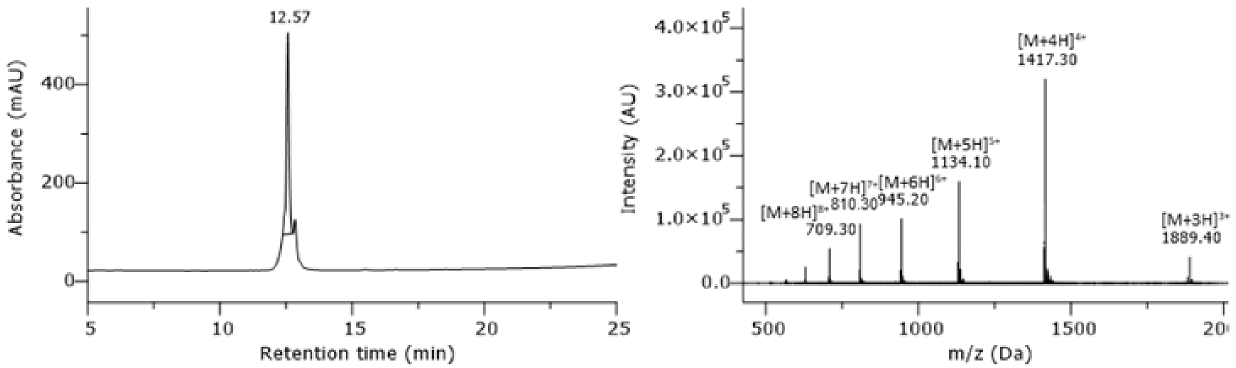

**Supplementary Fig. 12: HPLC chromatogram of purified peptide AMP #6.** Gradient 5-95% B in column 2, monitored at 220 nm.

$H_2N$-MYLKKFLALKNSLKKLSPFKCAVKSWLKKCAEVTFYSKLLGRRGKKDGN-$CONH_2$; peptide has been synthesized in 5 µmol scale. After purification (column 1, 05-50% MeCN), the 15 x TFA salt product (4.67 mg, 0.63 µmol, 13 %) was obtained as a white solid. $t_R$ = 12.57 min. Approximate purity > 90%. Formula: $C_{262}H_{433}N_{71}O_{62}S_3$. Molecular weight: 5665.87 g/mol. HRMS-ESI+ (m/z): $[M+6H]^{6+}$ calcd.: 945.2098; found: 945.2128.

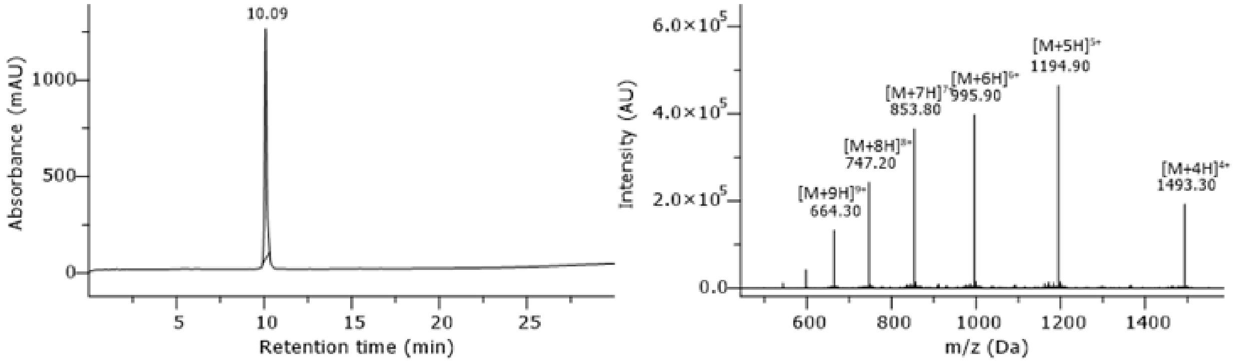

**Supplementary Fig. 13: HPLC chromatogram of purified peptide AMP #7.** Gradient 5-95% B in column 2, monitored at 220 nm.

$H_2N$-MRNFFKKTRLKYKGKKELIKKSRAFGLKTKKRSGFFFPRALKYEEEFY-$CONH_2$; peptide has been synthesized in 5 µmol scale. After purification (column 1, 05-50% MeCN), the 18 x TFA salt product (3.49 mg, 0.44 µmol, 9%) was obtained as a white solid. $t_R$ = 10.09 min. Purity ≥ 99%. Formula: $C_{282}H_{445}N_{77}O_{64}S$. Molecular weight: 5970.09 g/mol. HRMS-ESI+ (m/z): $[M+6H]^{6+}$ calcd.: 995.9029; found: 995.9010.

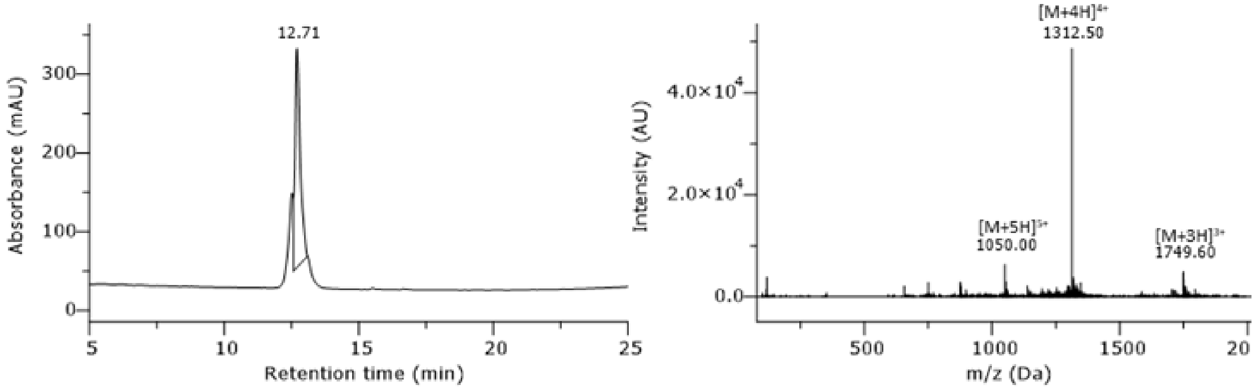

**Supplementary Fig. 14: HPLC chromatogram of purified peptide AMP #9.** Gradient 5-95% B in column 2, monitored at 220 nm.

$H_2N$-MGWWFPPKTGNGAGKAFKKAAAAKWGGLFLKAAWFANKGEWGGGFPKGY-$CONH_2$; peptide has been synthesized in 5 µmol scale. After purification (column 1, 05-50% MeCN), the 9 x TFA salt product (5.27 mg, 0.84 µmol, 17 %) was obtained as a white solid. $t_R$ = 12.71 min. Approximate purity 75%. Formula: $C_{255}H_{359}N_{65}O_{55}S$. Molecular weight: 5247.04 g/mol. HRMS-ESI+ (m/z): $[M+6H]^{6+}$ calcd.: 875.4589; found: 875.4609.

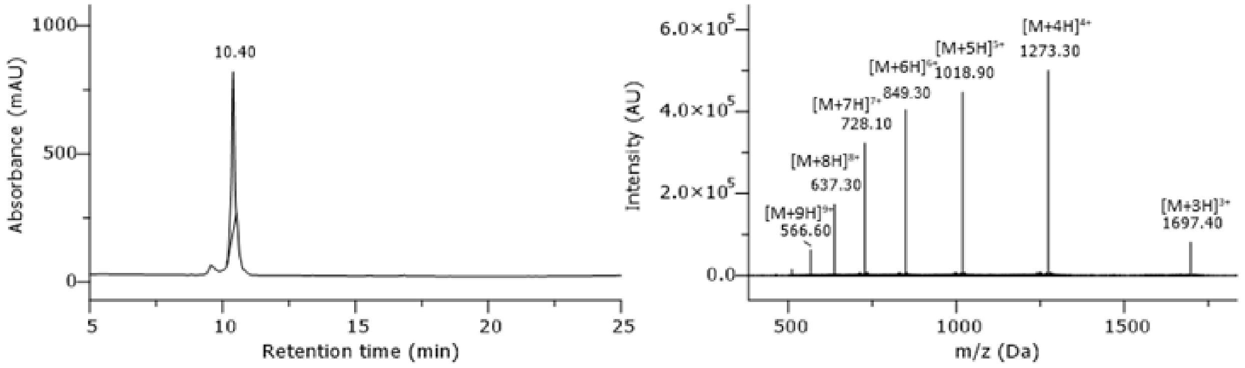

**Supplementary Fig. 15: HPLC chromatogram of purified peptide AMP #10.** Gradient 5-95% B in column 2, monitored at 220 nm.

$H_2N$-MCFCFKAGPKICRGLQRKKKKFKYQTSFTKTGFGFLTKPKSPAR-$CONH_2$; peptide has been synthesized in 5 µmol scale. After purification (column 1, 05-50% MeCN), the 14 x TFA salt product (2.98 mg, 0.45 µmol, 9%) was obtained as a white solid. $t_R$ = 10.40 min. Purity 93%. Formula: $C_{234}H_{376}N_{66}O_{53}S_4$. Molecular weight: 5090.15 g/mol. HRMS-ESI+ (m/z): $[M+6H]^{6+}$ calcd.: 849.3025; found: 849.3021.

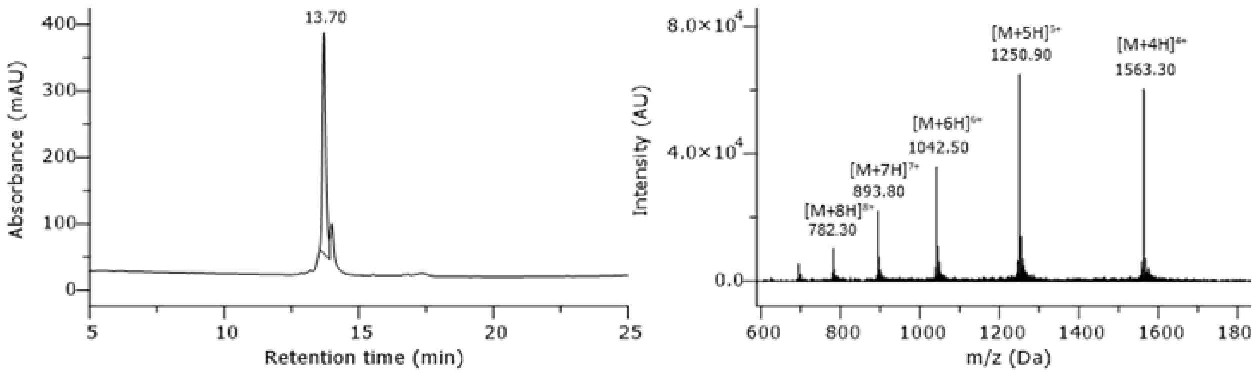

**Supplementary Fig. 16: HPLC chromatogram of purified peptide AMP #12.** Gradient 5-95% B in column 2, monitored at 220 nm.

$H_2N$-MWFDRKKFFWPGVCLFLLFFPKRFYKKGPEKVFSFRKKYKFARKCKCKL-$CONH_2$; peptide has been synthesized in 5 µmol scale. After purification (column 1, 05-75% MeCN), the 17 x TFA salt product (3.64 mg, 0.44 µmol, 9%) was obtained as a white solid. $t_R$ = 13.70 min. Approximate purity 86%. Formula: $C_{308}H_{458}N_{76}O_{56}S_4$. Molecular weight: 6249.66 g/mol. HRMS-ESI+ (m/z): $[M+9H]^{9+}$ calcd.: 695.3885; found: 695.3935.

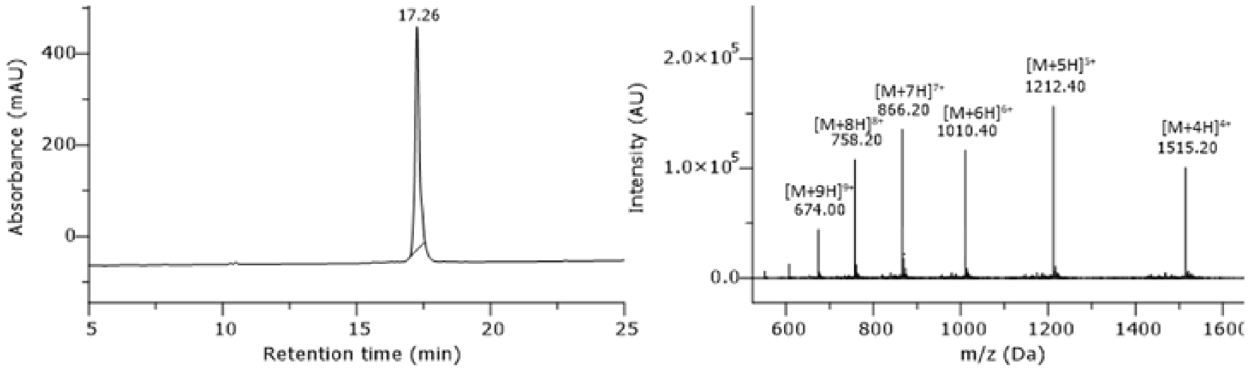

**Supplementary Fig. 17: HPLC chromatogram of purified peptide AMP #13**. Gradient 5-50% B, with addition of 1 mM TCEP, in column 2, monitored at 220 nm.

$H_2N$-MKQPSKTKTHFKYFLLFLKSVKKVAGFKKKKKKYHWRSYKEGSCFRKRT-$CONH_2$; peptide has been synthesized in 5 µmol scale. After purification (column 1, 05-50% MeCN), the 21 x TFA salt product (2.65 mg, 0.31 µmol, 6%) was obtained as a white solid. $t_R$ = 17.26 min. Purity ≥ 99%. Formula: $C_{285}H_{454}N_{80}O_{62}S_2$. Molecular weight: 6057.28 g/mol. HRMS-ESI+ (m/z): $[M+7H]^{7+}$ calcd.: 866.2165; found: 866.2123.

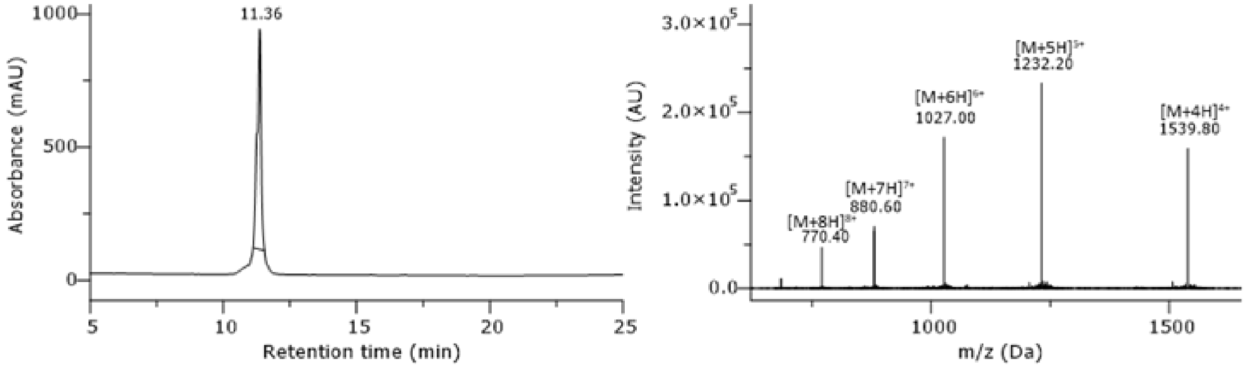

**Supplementary Fig. 18: HPLC chromatogram of purified peptide AMP #14.** Gradient 5-95% B in column 2, monitored at 220 nm.

$H_2N$-MKEKKFFFFCFKKRRGFYKRRFFCKTTCFTYCFYKPRGTKTMPYVFSE-$CONH_2$; peptide has been synthesized in 5 µmol scale. After purification (column 1, 05-50% MeCN), the 15 x TFA salt product (3.24 mg, 0.41 µmol, 8%) was obtained as a white solid. $t_R$ = 11.36 min. Approximate purity ≥ 89%. Formula: $C_{293}H_{427}N_{73}O_{62}S_6$. Molecular weight: 6156.36 g/mol. HRMS-ESI+ (m/z): $[M+7H]^{7+}$ calcd.: 880.5921; found: 880.5912.

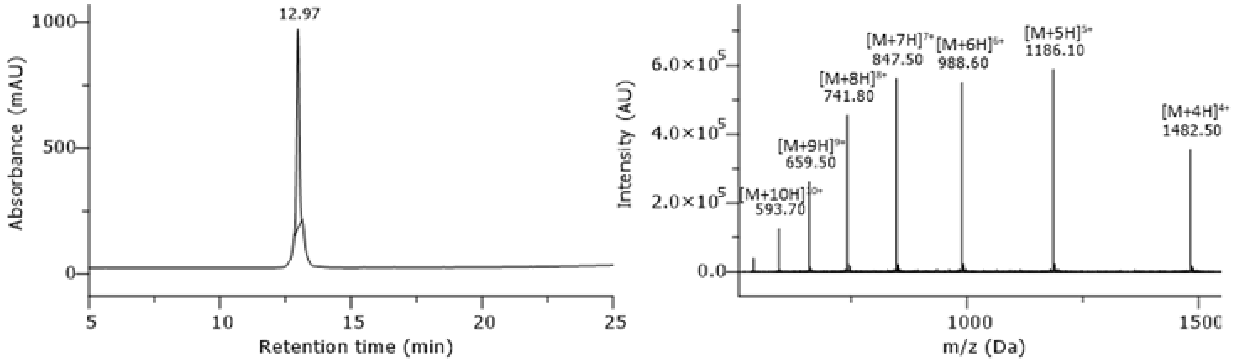

**Supplementary Fig. 19: HPLC chromatogram of purified peptide AMP #15.** Gradient 5-95% B in column 2, monitored at 220 nm.

$H_2N$-MEAFKKKPRLPLFKVKLTRFLERARGLGSYRIFEFFKKFGVKKFVSSLR-$CONH_2$; peptide has been synthesized in 5 µmol scale. After purification (column 1, 05-50% MeCN), the 16 x TFA salt product (3.69 mg, 0.48 µmol, 10 %) was obtained as a white solid. $t_R$ = 12.97 min. Purity ≥ 97%. Formula: $C_{283}H_{453}N_{77}O_{60}S$. Molecular weight: 5926.16 g/mol. HRMS-ESI+ (m/z): $[M+6H]^{6+}$ calcd.: 988.5834; found: 988.5846.

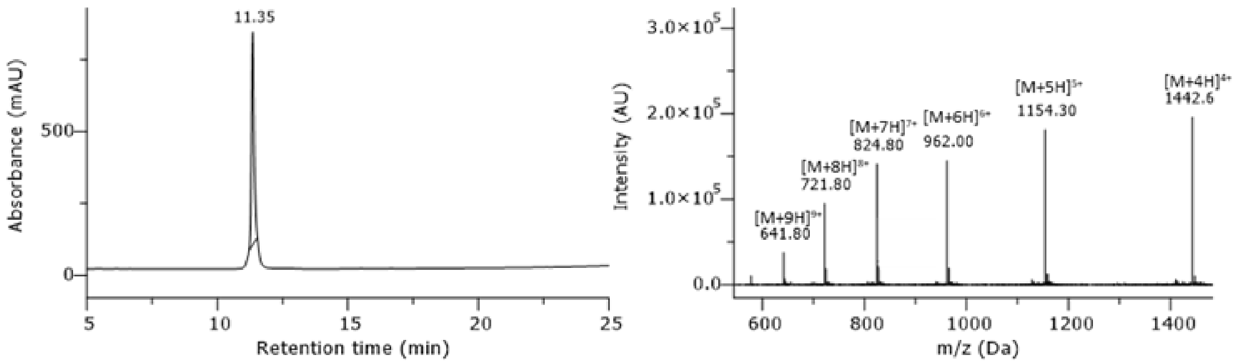

**Supplementary Fig. 20: HPLC chromatogram of purified peptide AMP #16.** Gradient 5-95% B in column 2, monitored at 220 nm.

$H_2N$-MKAKKWFESLFKTFFKKGKGIYPKSSFEKEKKTDKKKKFGGWVWFKK-$CONH_2$; peptide has been synthesized in 5 µmol scale. After purification (column 1, 05-50% MeCN), the 18 x TFA salt product (2.94 mg, 0.38 µmol, 8%) was obtained as a white solid. $t_R$ = 11.35 min. Purity ≥ 99%. Formula: $C_{281}H_{428}N_{68}O_{61}S$. Molecular weight: 5766.88 g/mol. HRMS-ESI+ (m/z): $[M+6H]^{6+}$ calcd.: 962.0454; found: 962.0462.

.

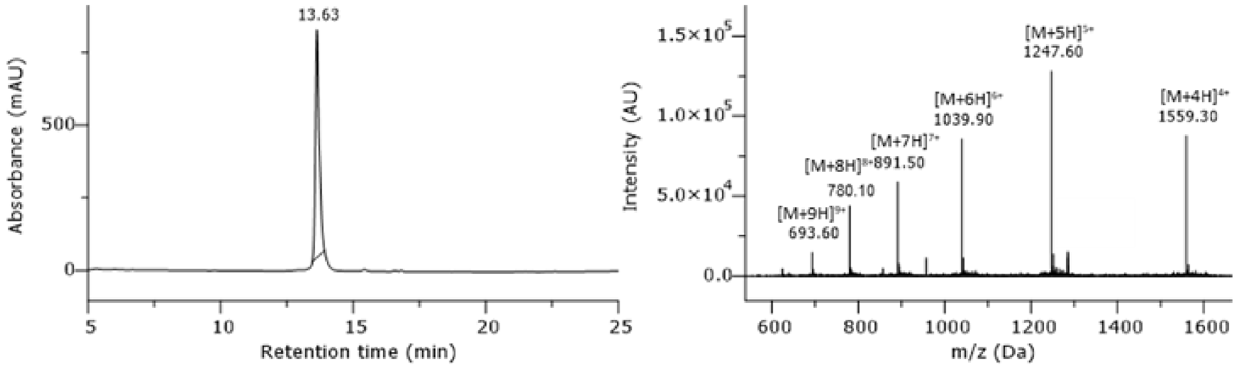

**Supplementary Fig. 21: HPLC chromatogram of purified peptide AMP #17.** Gradient 5-95% B in column 2, monitored at 220 nm.

$H_2N$-MWIKWKKPRKWGRRLKKKEKEELGDYIYLYCKVYRLFGFLPYFISKKTA-$CONH_2$; peptide has been synthesized in 5 µmol scale. After purification (column 1, 05-75% MeCN), the 16 x TFA salt product (2.73 mg, 0.34 µmol, 7 %) was obtained as a white solid. $t_R$ = 13.63 min. Purity ≥ 99%. Formula: $C_{301}H_{462}N_{76}O_{64}S_2$. Molecular weight: 6233.48 HRMS-ESI+ (m/z): $[M+7H]^{7+}$ calcd.: 891.3618; found: 891.3609.

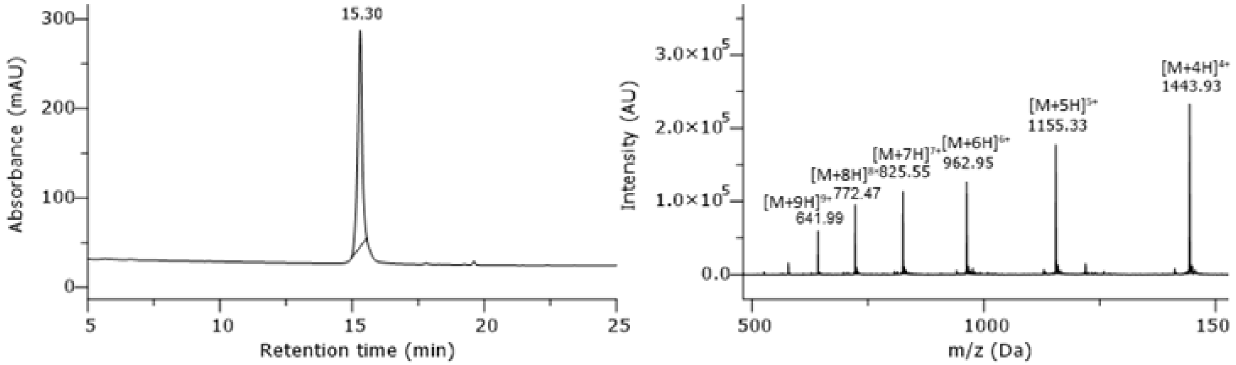

**Supplementary Fig. 22: HPLC chromatogram of purified peptide AMP #18.** Gradient 5-95% B in column 2, monitored at 220 nm.

$H_2N$-MCFFPRRKSKVKVKGGLCRLLFIFFKTTFCFKAKTKKEIKKGTGKKIVR-$CONH_2$; peptide has been synthesized in 5 µmol scale. After purification (column 1, 15-75% MeCN), the 17 x TFA salt product (2.99 mg, 0.39 µmol, 8%) was obtained as a white solid. $t_R$ = 15.30 min. Purity ≥ 99%. Formula: $C_{270}H_{451}N_{75}O_{56}S_4$. Molecular weight: 5772.19 HRMS-ESI+ (m/z): $[M+10H]^{10+}$ calcd.: 578.1443; found: 578.1465.

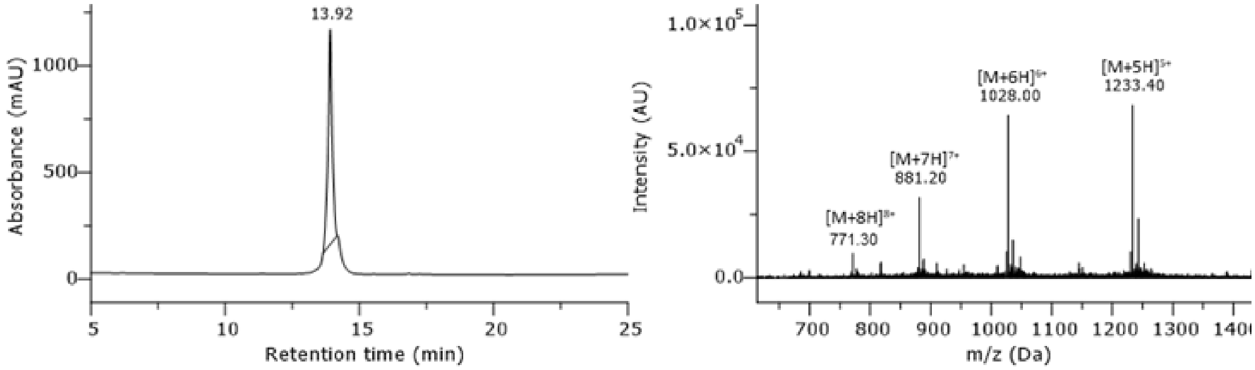

**Supplementary Fig. 23: HPLC chromatogram of purified peptide AMP #19.** Gradient 5-95% B in column 2, monitored at 220 nm.

$H_2N$-MCFCRYRFFYRRRIRFFKWGPYFYVWFGFPFGRKAFFLSVFRRRFC-$CONH_2$; peptide has been synthesized in 5 µmol scale. After purification (column 1, 05-75% MeCN), the 13 x TFA salt product (2.24 mg, 0.29 µmol, 6 %) was obtained as a white solid. $t_R$ = 13.92 min. Approximate purity > 90%. Formula: $C_{305}H_{417}N_{81}O_{51}S_4$. Molecular weight: 6162.34 g/mol. HRMS-ESI+ (m/z): $[M+6H]^{6+}$ calcd.: 1028.0324; found: 1028.0393.

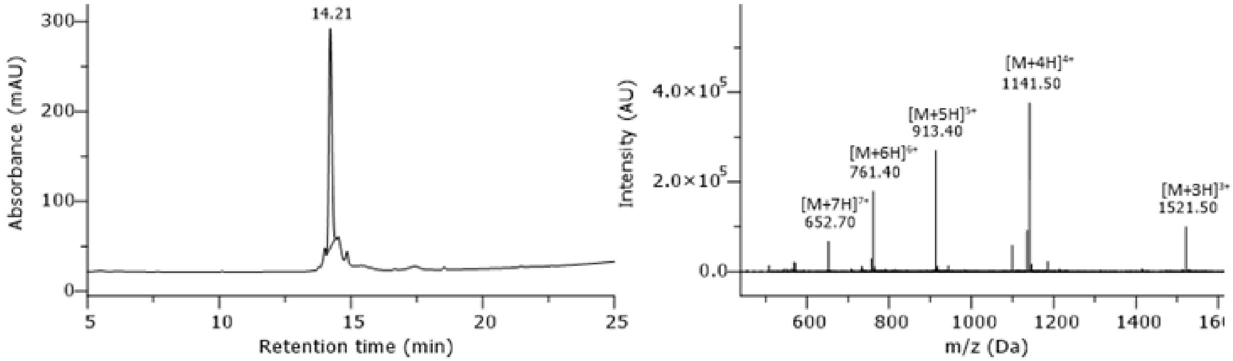

**Supplementary Fig. 24: HPLC chromatogram of purified peptide AMP #20.** Gradient 5-95% B in column 2, monitored at 220 nm.

$H_2N$-MRGRPPKRIRSVIIAQTTTATAKKIVIVLLLIFSSSKRRR-$CONH_2$; peptide has been synthesized in 5 µmol scale. After purification (column 1, 15-75% MeCN), the 12 x TFA salt product (0.35 mg, 0.06 µmol, 1 %) was obtained as a white solid. $t_R$ = 14.21 min. Approximate purity > 85%. Formula: $C_{203}H_{367}N_{67}O_{49}S$. Molecular weight: 4562.57 g/mol. HRMS-ESI+ (m/z): $[M+5H]^{5+}$ calcd.: 913.3683; found: 913.3685.

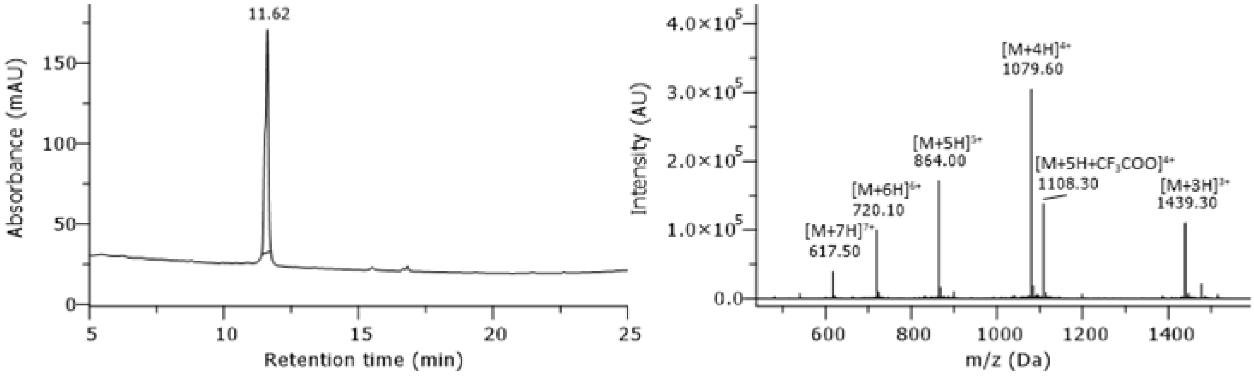

**Supplementary Fig. 25: HPLC chromatogram of purified peptide AMP #21.** Gradient 5-95% B in column 2, monitored at 220 nm.

$H_2N$-MGIGKFQKMRFIGAIRASKGVAKGLLRIAAIRTGRRALTT-$CONH_2$; peptide has been synthesized in 5 µmol scale. After purification (column 1, 05-50% MeCN), the 11 x TFA salt product (2.50 mg, 0.45 µmol, 9 %) was obtained as a white solid. $t_R$ = 11.62 min. Approximate purity > 68%. Formula: $C_{191}H_{338}N_{64}O_{45}S_2$. Molecular weight: 4315.25 g/mol. HRMS-ESI+ (m/z): $[M+7H]^{7+}$ calcd.: 617.3733; found: 617.3767.

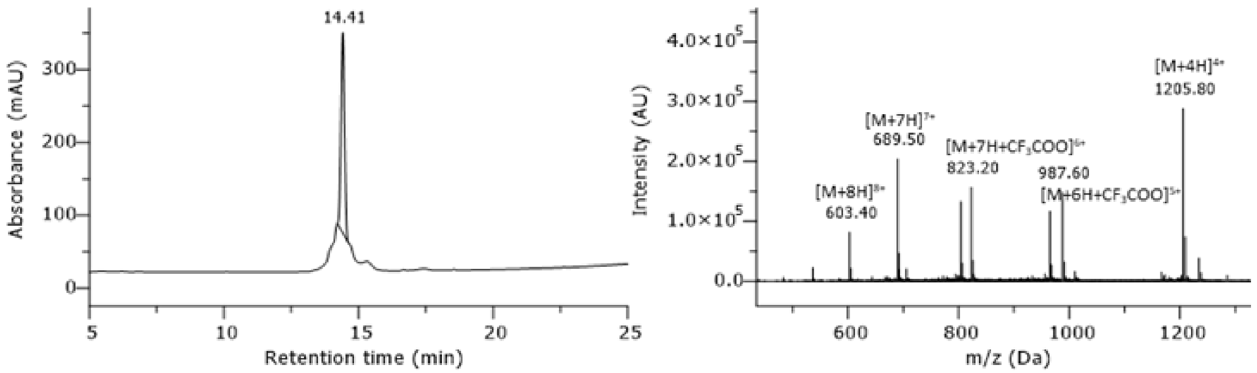

**Supplementary Fig. 26: HPLC chromatogram of purified peptide AMP #23.** Gradient 5-95% B in column 2, monitored at 220 nm.

$H_2N$-MSTRSSSIRRLVEAVRTRFRAALRTVLFFALRTTKRRPRR-$CONH_2$; peptide has been synthesized in 5 µmol scale. After purification (column 1, 15-75% MeCN), the 14 x TFA salt product (2.01 mg, 0.31 µmol, 6%) was obtained as a white solid. $t_R$ = 14.41 min. Approximate purity 84%. Formula: $C_{209}H_{366}N_{78}O_{51}S$. Molecular weight: 4819.69 g/mol. HRMS-ESI+ (m/z): $[M+5H]^{5+}$ calcd.: 964.7716; found: 964.7743.

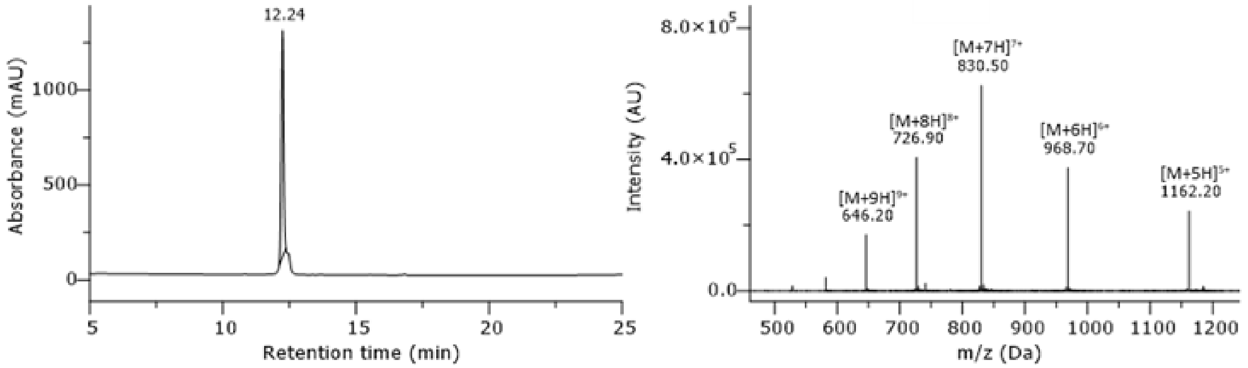

**Supplementary Fig. 27: HPLC chromatogram of purified peptide AMP #24.** Gradient 5-95% B in column 2, monitored at 220 nm.

$H_2N$-MAIRIIGRLARVRARVVARVRSRLLADDPPEDLLRVARRRKGRRWLFLS-$CONH_2$; peptide has been synthesized in 5 µmol scale. After purification (column 1, 05-50% MeCN), the 16 x TFA salt product (5.74 mg, 0.75 µmol, 15 %) was obtained as a white solid. $t_R$ = 12.24 min. Approximate purity > 97%. Formula: $C_{255}H_{448}N_{94}O_{59}S$. Molecular weight: 5806.94 g/mol. HRMS-ESI+ (m/z): $[M+6H]^{6+}$ calcd.: 968.7530; found: 968.7544.

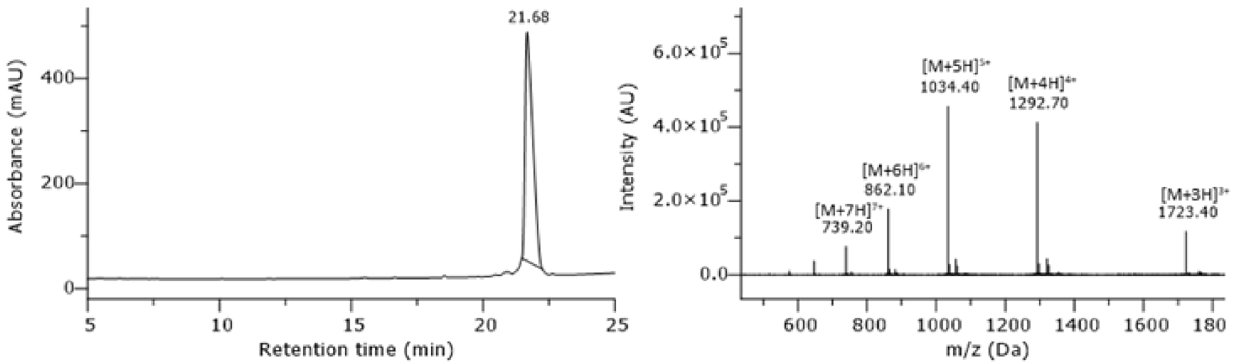

**Supplementary Fig. 28: HPLC chromatogram of purified peptide AMP #26.** Gradient 5-95% B in column 2, monitored at 220 nm.

$H_2N$-MKLRRRLRTRMVALLVLGVLFLLMLFFIIFLRRMLMRRFRA-$CONH_2$; peptide has been synthesized in 5 µmol scale. After purification (column 1, 20-95% MeCN), the 12 x TFA salt product (4.25 mg, 0.65 µmol, 13 %) was obtained as a white solid. $t_R$ = 21.68 min. Purity 96%. Formula: $C_{241}H_{415}N_{73}O_{42}S_5$. Molecular weight: 5167.66 g/mol. HRMS-ESI+ (m/z): $[M+5H]^{5+}$ calcd.: 1034.4324; found: 1034.4339.

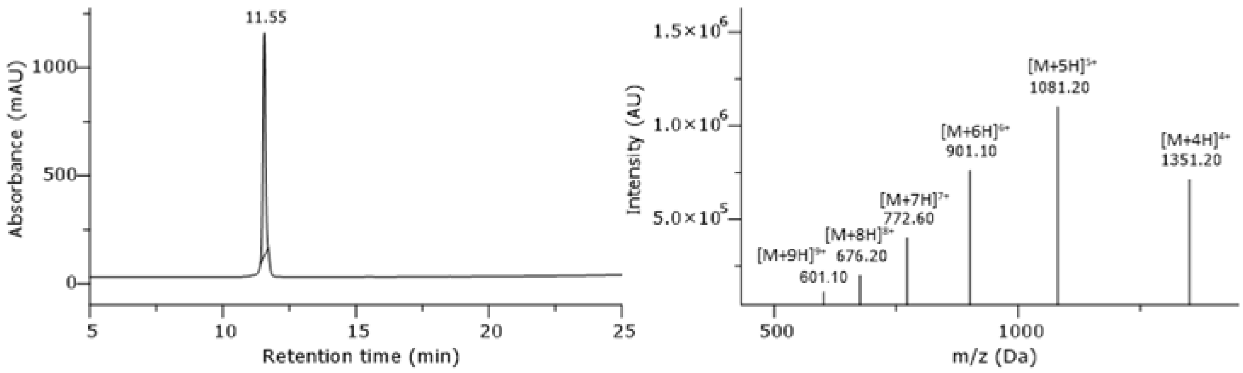

**Supplementary Fig. 29: HPLC chromatogram of purified peptide AMP #27.** Gradient 5-95% B in column 2, monitored at 220 nm.

$H_2N$-MNTTSNMIHRAVQQKRISFRAAKLTVLFLFKRRLLRRLLRHHEN-$CONH_2$; peptide has been synthesized in 5 µmol scale. After purification (column 1, 05-50% MeCN), the 15 x TFA salt product (4.03 mg, 0.57 µmol, 11%) was obtained as a white solid. $t_R$ = 11.55 min. Purity ≥ 99%. Formula: $C_{239}H_{405}N_{83}O_{56}S_2$. Molecular weight: 5401.42 g/mol. HRMS-ESI+ (m/z): $[M+5H]^{5+}$ calcd.: 1081.225; found: 1081.2283.

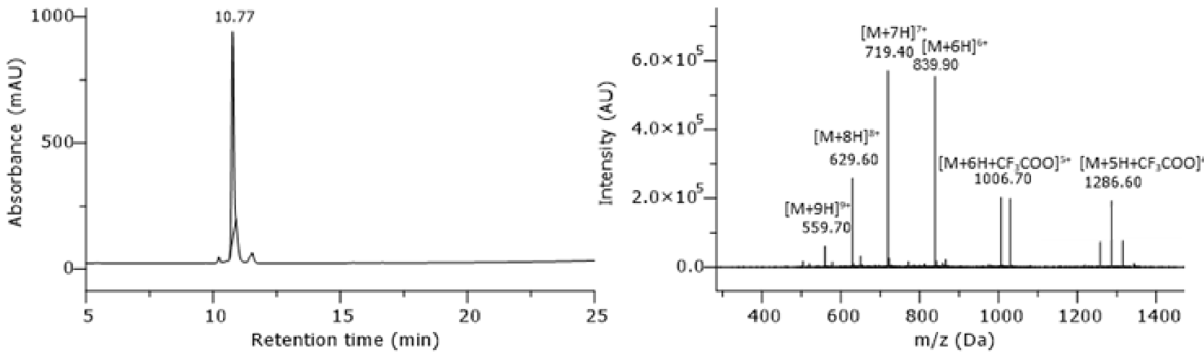

**Supplementary Fig. 30: HPLC chromatogram of purified peptide AMP #28.** Gradient 5-95% B in column 2, monitored at 220 nm.

$H_2N$-MMKIRNTLRSRKEAVRRIFSLRRRSVFTEMARAFRRFKAR-$CONH_2$; peptide has been synthesized in 5 µmol scale. After purification (column 1, 05-50% MeCN), the 16 x TFA salt product (4.53 mg, 0.66 µmol, 13 %) was obtained as a white solid. $t_R$ = 10.77 min. Purity 90%. Formula: $C_{218}H_{377}N_{81}O_{50}S_3$. Molecular weight: 5029.03 g/mol. HRMS-ESI+ (m/z): $[M+7H]^{7+}$ calcd.: 719.4170; found: 719.4181.

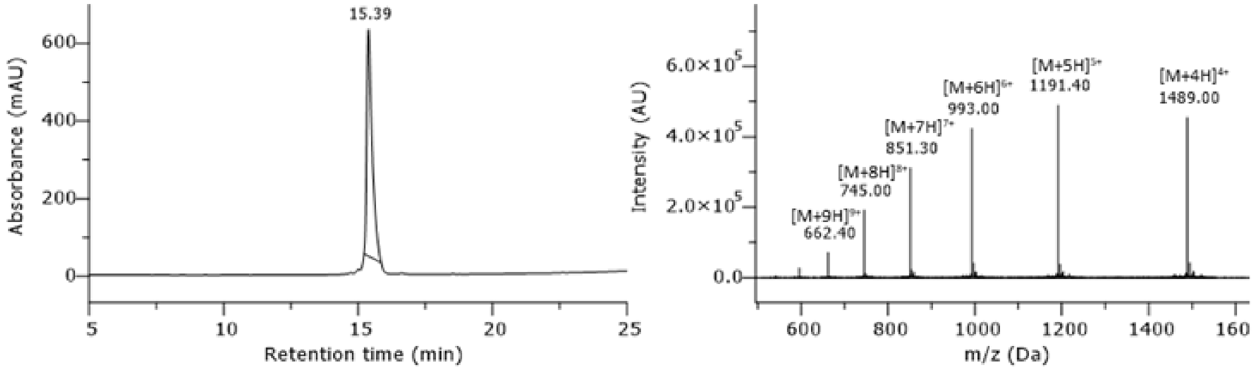

**Supplementary Fig. 31: HPLC chromatogram of purified peptide AMP #29.** Gradient 5-95% B in column 2, monitored at 220 nm.

$H_2N$-MKKKKKGILKQNNKKKKYTLFNRMVVFLFLGFIMIIFVQKYKKVIYHK-$CONH_2$; peptide has been synthesized in 5 µmol scale. After purification (column 1, 05-75% MeCN), the 17 x TFA salt product (4.80 mg, 0.61 µmol, 12 %) was obtained as a white solid. $t_R$ = 15.39 min. Purity ≥ 99%. Formula: $C_{287}H_{471}N_{73}O_{57}S_3$. Molecular weight: 5952.45 g/mol. HRMS-ESI+ (m/z): $[M+5H]^{5+}$ calcd.: 1191.3161; found: 1191.3205.

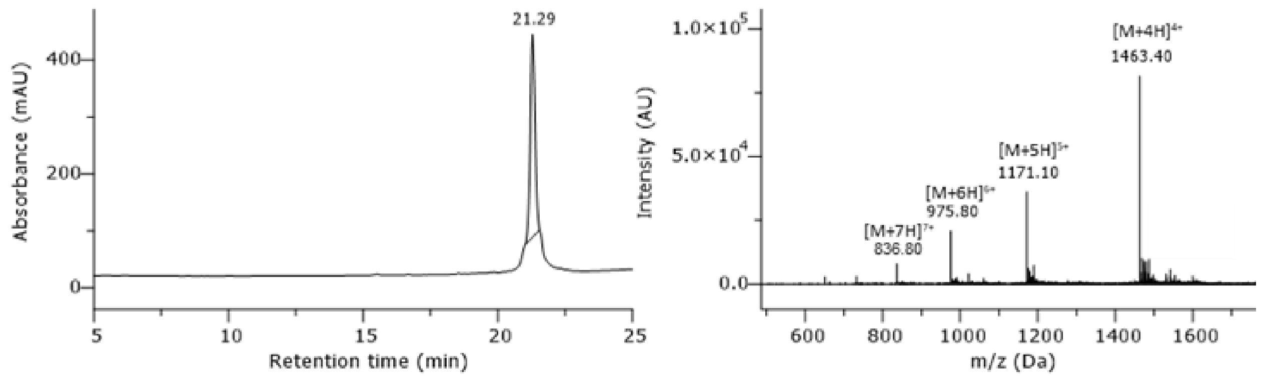

**Supplementary Fig. 32: HPLC chromatogram of purified peptide AMP #30.** Gradient 5-95% B in column 2, monitored at 220 nm.

$H_2N$-MGFGLWGLFHFKMNVPNLFKNGFIFLIIMIFTVWGLFFGKKKAYIEKFL-$CONH_2$; peptide has been synthesized in 5 µmol scale. After purification (column 1, 20-95% MeCN), the 8 x TFA salt product (1.44 mg, 0.21 µmol, 4 %) was obtained as a white solid. $t_R$ = 21.29 min. Approximate purity 98%. Formula: $C_{295}H_{429}N_{63}O_{56}S_3$**.** Molecular weight: 5850.14 g/mol. HRMS-ESI+ (m/z): $[M+5H]^{5+}$ cacld.: 1170.8453; found: 1170.8442.

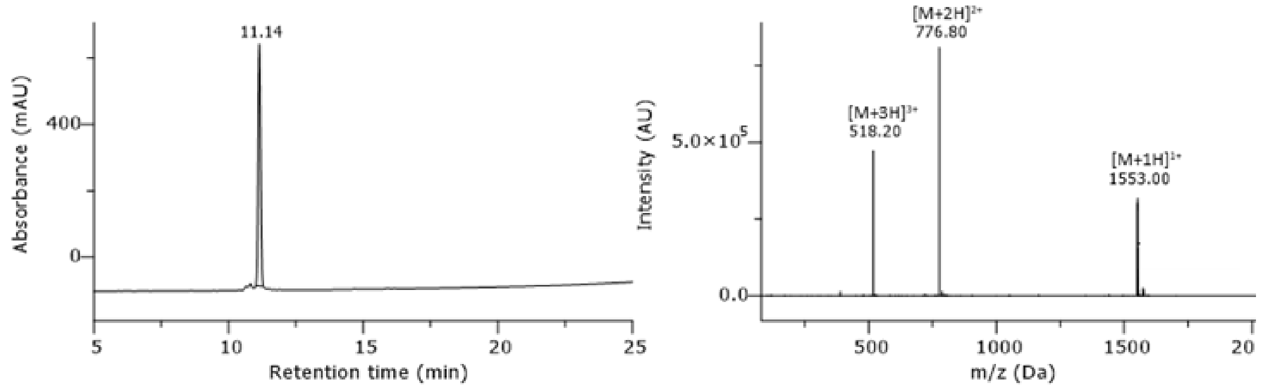

**Supplementary Fig. 33 HPLC chromatogram of purified peptide BP100.** Gradient 5-95% B in column 2, monitored at 220 nm.

$H_2N$-MKKLFKKILKYL-$CONH_2$; peptide has been synthesized in 5 µmol scale. After purification (column 1, 05-50% MeCN), the 6 x TFA salt product (4.03 mg, 1.80 µmol, 36 %) was obtained as a white solid. $t_R$ = 11.14 min. Purity 97%. Formula: $C_{77}H_{134}N_{18}O_{13}S$. Molecular weight: 1552.06 g/mol. HRMS-ESI+ (m/z): $[M+2H]^{2+}$ calcd.: 776.5122; found: 776.5110.

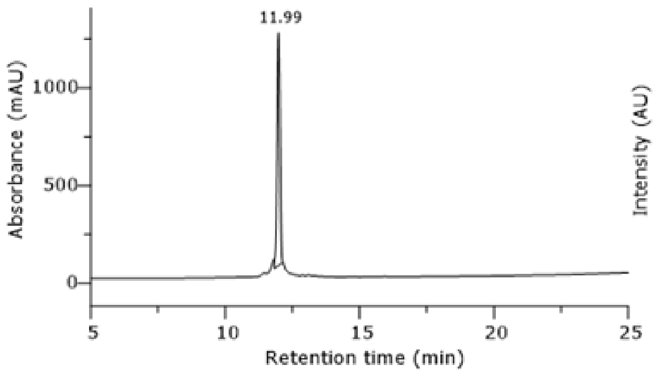

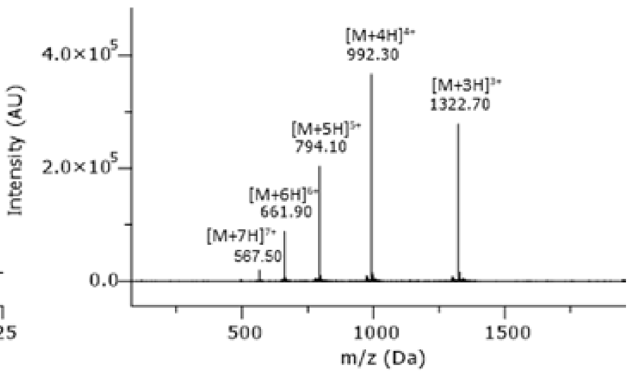

**Supplementary Fig. 34: HPLC chromatogram of purified peptide Cecropin B.** Gradient 5-95% B in column 2, monitored at 220 nm.

$H_2N$-MKWKVFKKIEKMGRNIRNGIVKAGPAIAVLGEAKAL-$CONH_2$; peptide has been synthesized in 5 µmol scale. After purification (column 1, 05-50% MeCN), the 10 x TFA salt product (2.29 mg, 0.45 µmol, 9%) was obtained as a white solid. $t_R$ = 11.99 min. Purity 97%. Formula: $C_{181}H_{311}N_{53}O_{42}S_2$. Molecular weight: 3965.86 g/mol. HRMS-ESI+ (m/z): $[M+4H]^{4+}$ calcd.: 992.3404; found: 992.3409.

#### Supplementary references:

1. Espah Borujeni, A. & Salis, H. M. Translation initiation is controlled by RNA folding kinetics via a ribosome drafting mechanism. *J. Am. Chem. Soc.* **138**, 7016–7023 (2016).
2. Salis, H. M., Mirsky, E. A. & Voigt, C. A. Automated design of synthetic ribosome binding sites to control protein expression. *Nat. Biotechnol.* **27**, 946–950 (2009).
3. Das, P. *et al.* Accelerated antimicrobial discovery via deep generative models and molecular dynamics simulations. *Nature Biomedical Engineering* **5**, 613–623 (2021).
4. Jumper, J. *et al.* Highly accurate protein structure prediction with AlphaFold. *Nature* **596**, 583–589 (2021).
5. Tucs, A. *et al.* Generating Ampicillin-Level Antimicrobial Peptides with Activity-Aware Generative Adversarial Networks. *ACS Omega* **5**, 22847–22851 (2020).
6. Flamm, C., Fontana, W., Hofacker, I. L. & Schuster, P. RNA folding at elementary step resolution. *RNA* **6**, 325–338 (2000).
7. Cock, P. J. A. *et al.* Biopython: freely available Python tools for computational molecular biology and bioinformatics. *Bioinformatics* **25**, 1422–1423 (2009).
8. Müller, A. T., Gabernet, G., Hiss, J. A. & Schneider, G. modlAMP: Python for antimicrobial peptides. *Bioinformatics* **33**, 2753–2755 (2017).
